# CucurbitAgent: An AI agent for exploring cucurbit multi-omics resources

**DOI:** 10.64898/2026.10.09.758138

**Authors:** Rong Zhang, Jian Zhang, Yuming Zhang, Xiaofei Zhang, Jingjing Yang, Changxuan Xia, Jialong Zhang, Hong Zhao, Changlong Wen

**Affiliations:** National Engineering Research Center for Vegetables; State Key Laboratory of Vegetable Biobreeding, Beijing Vegetable Research Center, Beijing Academy of Agriculture and Forestry Sciences, Beijing 100097, China; Key Laboratory of Biology and Genetic Improvement of Horticultural Crops (North China); Key Laboratory of Superior Quality Vegetable Germplasm Innovation, Ministry of Agriculture and Rural Affairs; Key Laboratory of Urban Agriculture (North), Ministry of Agriculture and Rural Affairs, Beijing Vegetable Research Center, Beijing Academy of Agriculture and Forestry Sciences, Beijing 100097, China; Beijing Key Laboratory of Vegetable Germplasms Improvement; Beijing Key Laboratory of Molecular and Intelligent Breeding for Crops, Beijing 100097, China; MOE Key Laboratory of Bioinformatics, State Key Laboratory of Molecular Oncology, Beijing Frontier Research Center for Biological Structure, School of Pharmaceutical Sciences, Tsinghua University, Beijing 100084, China

**Keywords:** AI agent, large language model, functional genomics, Cucurbitaceae, multi-omics integration

## Abstract

Functional genomics in crops increasingly depends on integrating information from an extensive literature and heterogeneous multi-omics datasets. Large language models (LLMs) offer a powerful approach to synthesizing these resources and revealing hidden connections across datasets; however, they frequently hallucinate, producing plausible but unsupported claims, when responding to gene-specific queries. Here, we present CucurbitAgent, an AI research agent for functional genomics in the Cucurbitaceae, a major family of horticultural crops. CucurbitAgent integrates multi-omics resources from nine cultivated cucurbit species and more than 18,000 publications, encompassing approximately 256,000 genes, tissue-resolved expression profiles, over 16 million natural genetic variants, and roughly 242,000 predicted protein structures with more than 14 million predicted binding-site residue annotations. Responses link retrieved evidence to source records, including gene entries, expression measurements, structural predictions, and literature passages. On a curated benchmark of 50 tasks assessed by domain experts under blinded conditions, CucurbitAgent achieved an overall pairwise win rate of 96.0% against nine general-purpose LLMs, based on weighted scores for factual accuracy, completeness, relevance, and traceability. The platform is freely available at https://cucurbitagent.com and supports queries in 37 languages collectively spoken by an estimated 70% or more of the world’s population. CucurbitAgent not only unifies evidence retrieval and interpretation but also reveals latent associations across datasets, helping researchers formulate testable hypotheses and substantially accelerating candidate-gene research. Its framework is readily transferable to other crops and, more broadly, provides a blueprint for building specialized agents in any domain.

## INTRODUCTION

Research in crop functional genomics increasingly depends on integrating information from an extensive literature and heterogeneous multi-omics datasets (Depuydt et al., 2023; Stark et al., 2019). These resources are distributed across specialized databases and an expanding publication record. Manually querying these scattered resources is time-consuming and makes it difficult to integrate the evidence needed to formulate hypotheses (Tarazona et al., 2021). Large language models (LLMs) offer a way to synthesize this fragmented information (Lewis et al., 2020). However, they can generate plausible but incorrect statements (Farquhar et al., 2024), a concern for queries that require precise gene or tissue information. Such limitations are especially consequential for crop molecular breeding, a field that relies heavily on integrating specialized, species-specific knowledge.

AI assistants tailored to plant research are emerging. PlantScience.ai serves the broader plant science community (Yu et al., 2026), and PlantGPT supports functional-genomics queries in Arabidopsis (Provart et al., 2021; Zhang et al., 2025). Crop-specific resources include SeedLLM·Rice, which couples a rice-specific language model with a biological knowledge graph (Yang et al., 2025), and the Potato Knowledge Hub, which integrates literature retrieval with functional gene records and analytical tools (Li et al., 2026). Biomedical models such as BioGPT (Luo et al., 2022) and BioMedLM (Bolton et al., 2024) further illustrate approaches to domain adaptation. However, such assistants remain unavailable for the Cucurbitaceae, a major horticultural family (Grumet et al., 2021) that includes cucumber, watermelon, melon, and other cultivated crops. As genomic resources and the associated literature for this family continue to expand (Marks et al., 2021; Sun et al., 2022), coexisting reference assemblies within individual species further complicate identifier resolution (Guan et al., 2024; Li et al., 2019). These considerations motivate the development of an AI-driven research assistant that reconciles gene identifiers and integrates multi-omics data across multiple cucurbit species, thereby accelerating functional genomics research in this family.

Here, we present CucurbitAgent, an LLM-based research agent that coordinates specialized tools to retrieve and integrate cucurbit multi-omics evidence. The system has three layers: a top-level tool-calling agent that selects tools and synthesizes results, four independent evidence modules— Gene, Expression, Protein, and Literature—and a knowledge-foundation layer containing the underlying databases. Its central design principle is to make retrieved evidence traceable to source records, so that users can inspect the basis of any answer. The knowledge foundation integrates multi-omics data from nine cultivated cucurbit species, spanning approximately 256,000 genes, tissue-resolved expression profiles, and more than 16 million natural variants. It also includes roughly 242,000 protein structures predicted by ESMFold (Lin et al., 2023), over 14 million predicted binding-site residue annotations, and a curated corpus of over 18,000 publications on cucurbit functional genomics.

To broaden access, the platform supports queries in 37 languages reaching more than 70% of the global population (Eberhard et al., 2025). We evaluated CucurbitAgent on an expert-authored benchmark of 50 tasks under blinded expert review. The system achieved an overall pairwise win rate of 96.0% against nine general-purpose LLMs based on weighted composite scores. Expert evaluation indicated strong performance in factual accuracy and traceability on the benchmark tasks, supporting its use for retrieving and integrating cucurbit research evidence. By linking answers to underlying records, the platform aims to facilitate candidate-gene analysis and hypothesis development. The platform is freely available at https://cucurbitagent.com. Beyond the cucurbits, its design offers a general template for building trustworthy research assistants in other crops and specialized domains.

## RESULTS

### Architecture and design of CucurbitAgent

CucurbitAgent was designed to help researchers integrate heterogeneous evidence through a single natural-language interface and uncover relationships across data types. The system has three layers (Figure 1): a top-level tool-calling agent, four independent evidence modules, and a knowledge-foundation layer containing the four underlying databases. Its central design principle is to make retrieved evidence traceable to source records. The four modules—Gene, Expression, Protein, and Literature—provide gene identifiers and functional annotations, tissue expression profiles and co-expression neighbors, predicted protein structures and candidate binding sites, and specialist literature, respectively. These resources represent complementary types of evidence for candidate-gene analysis. The agent is based on a large language model operating in a multi-turn reasoning-and-acting (ReAct) loop (Yao et al., 2023). Given a question, it calls the relevant modules, integrates the retrieved evidence, and returns an answer linked to the underlying records. Tool calls, retrieved evidence, and progress updates are displayed to users in real time. LLM hallucinations are a concern in scientific queries (Farquhar et al., 2024), including queries requiring precise identifiers, coordinates, or citations. Retrieval provides a means of grounding responses in external records (Lewis et al., 2020), although it does not guarantee that every generated statement is correct. Each module also operates independently: researchers can search it directly and refine the results through multi-turn dialogue (Supplemental Figures 1–3). Alternatively, they can use the top-level agent to retrieve evidence across modules. A module inventory is provided in Supplemental Table 1. To support researchers working in languages other than English, the interface provides text interaction in 37 languages (full list in Supplemental Table 2), including the ten most widely spoken languages worldwide; together, these languages are used by over 70% of people worldwide (Eberhard et al., 2025). Answers are returned in the user’s language, while source evidence, gene identifiers, and citations are preserved (implementation details in Methods).

**Figure 1.**
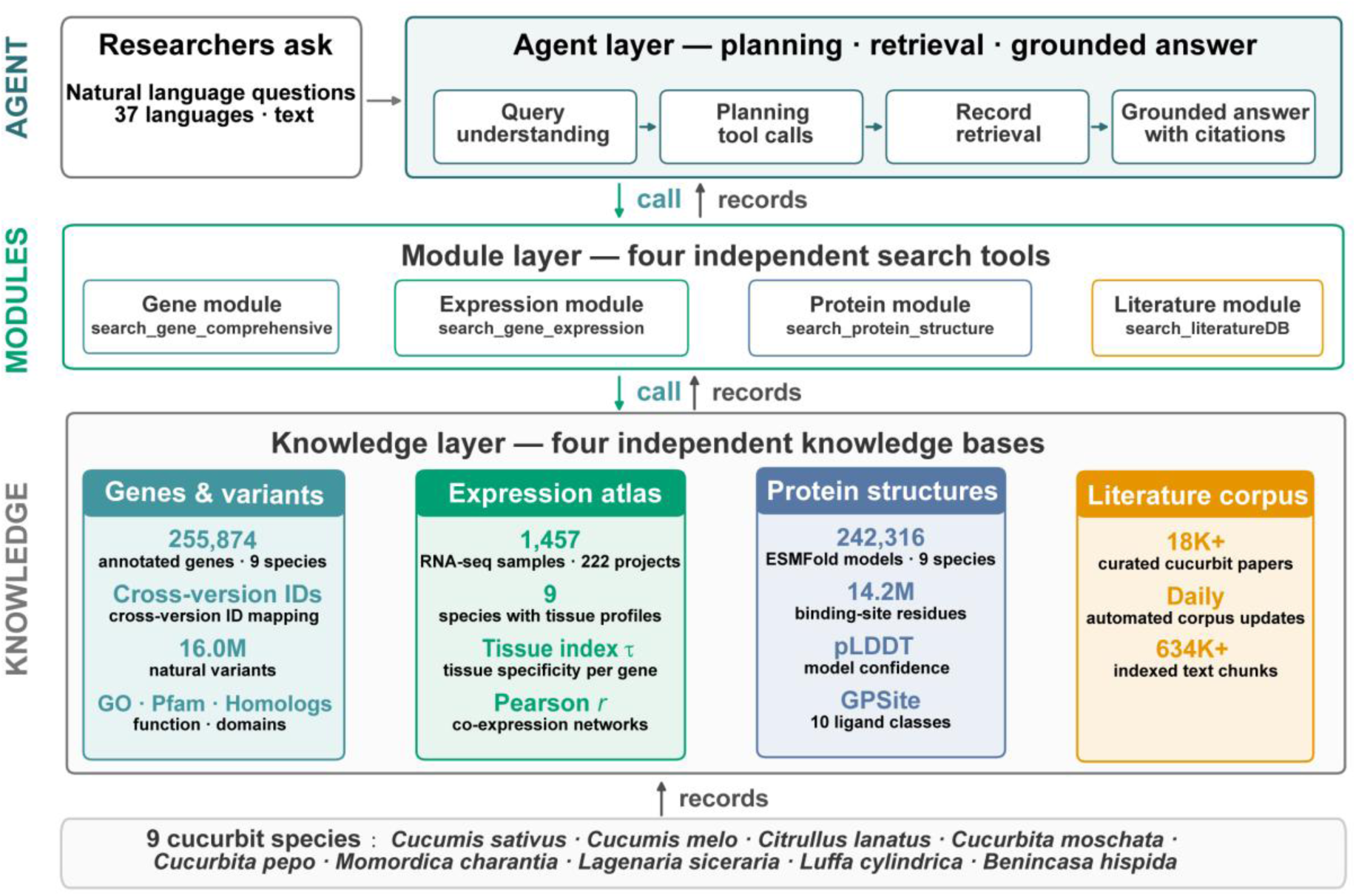
Three-layer architecture of CucurbitAgent. Researchers submit natural-language questions through an interface supporting 37 languages. The agent layer processes questions through an iterative sequence of query interpretation, tool selection, record retrieval, and evidence-grounded answer generation with citations. The module layer comprises four independent modules—Gene, Expression, Protein, and Literature—each available to the agent as a search tool and to researchers as a standalone search interface. The knowledge-foundation layer hosts four databases—Genes and Variants, Expression Atlas, Protein Structures, and Literature Corpus—built from multi-omics data and literature spanning nine cucurbit species.

### A pan-cucurbit multi-omics knowledge foundation

The knowledge foundation comprises three multi-omics modules—Gene, Expression, and Protein. The Gene module provides a unified entry point for gene-level information across the nine cultivated species. It integrates resources for cucumber (Cucumis sativus), melon (C. melo), watermelon (Citrullus lanatus), pumpkin and squash (Cucurbita moschata and C. pepo), bitter gourd (Momordica charantia), bottle gourd (Lagenaria siceraria), sponge gourd (Luffa cylindrica), and wax gourd (Benincasa hispida), totaling 255,874 genes (Figure 2A, per-species inventory in Supplemental Table 3). The module contains reference gene structures and protein sequences, Gene Ontology (GO) annotations (Gene Ontology Consortium, 2021), Pfam domain annotations (Mistry et al., 2021), and candidate homologs (identity ≥ 40%, coverage ≥ 50%) in the Cucurbitaceae and in Arabidopsis, maize, rice, and tomato. Explicit cross-version maps cover the four species with multiple reference assemblies (Supplemental Table 3)—cucumber ChineseLong V1–V3, watermelon 97103 V1–V2.5, melon DHL92 V3.5.1–V4.0, and C. moschata Rifu V1–V2— so that any legacy identifier from older literature resolves to a unified gene card. Natural-variant data currently cover five species—cucumber, watermelon, melon, pumpkin (C. moschata), and squash (C. pepo)—with 16.0 million indexed sites in total (Figure 2B). Each variant is classified as coding or non-coding based on its genomic location. Coding variants are further labeled as synonymous or protein-altering, and for protein-altering variants, the resulting amino-acid change is recorded. On each gene card, variants are summarized by count and consequence class, and CDS and protein sequences of any gene can be downloaded directly.

**Figure 2.**
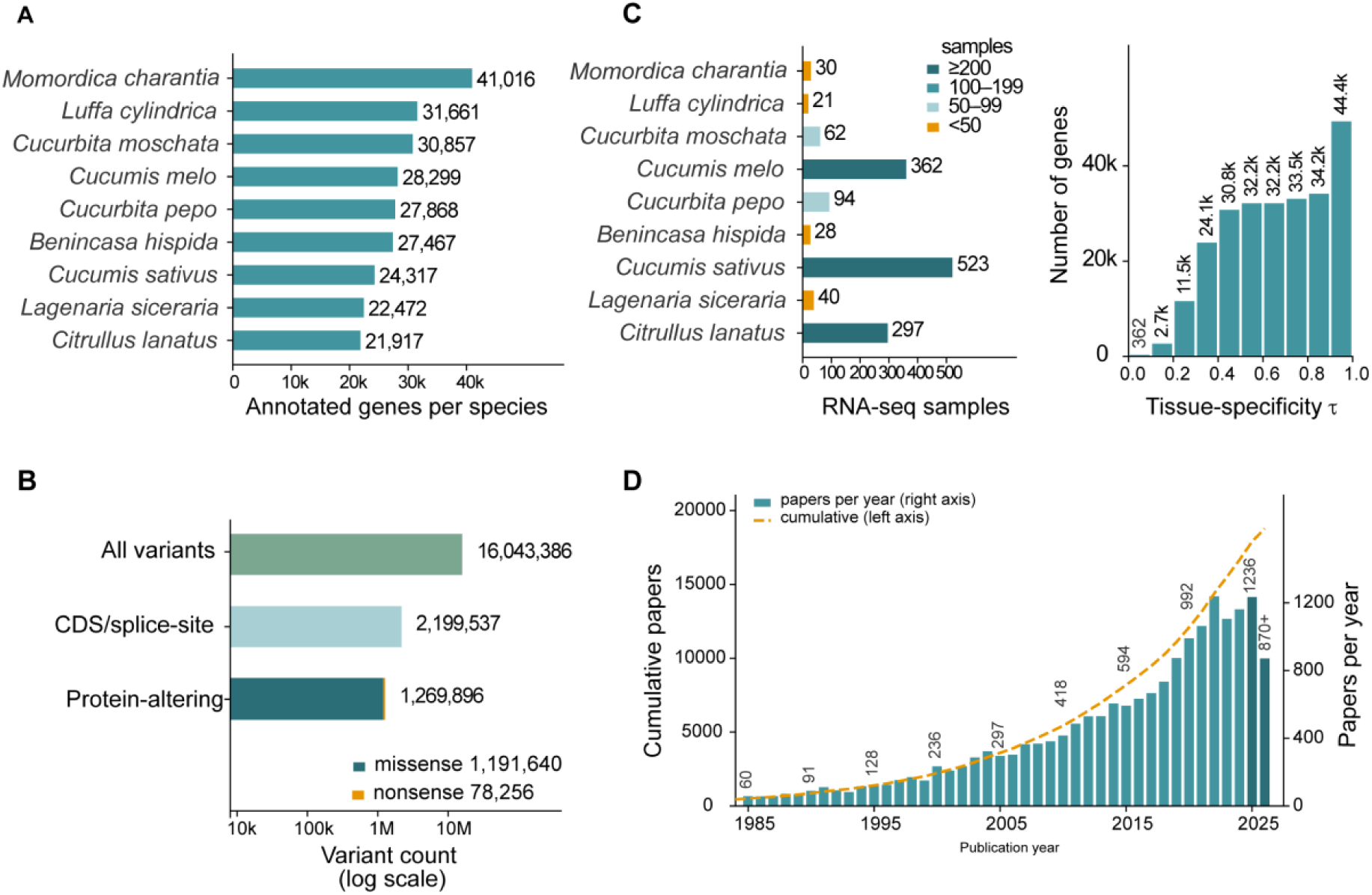
Composition and scale of the pan-cucurbit knowledge foundation. (A) Number of annotated genes in the reference annotation of each of the nine species. Cross-version identifiers were reconciled for the four species with multiple supported assemblies. (B) Variant annotation funnel (logarithmic x axis): 16,043,386 indexed variants from five species, including 2,199,537 within coding sequences or splice sites and 1,269,896 predicted to alter protein sequences (1,191,640 missense and 78,256 nonsense variants). (C) Left, number of RNA-seq samples per species, colored by sample-size category; right, distribution of the tissue-specificity index (tau). (D) Papers in the literature corpus by publication year (bars, right axis) and cumulative paper count (orange dashed curve, left axis). The September 8, 2026 snapshot includes 18,305 papers retained after filtering. The final bar (870+) covers part of 2026.

The Expression module consolidates public RNA-seq resources for the nine species into a transcriptome atlas of 1,457 samples, indexed by species, project, and tissue (Figure 2C). The atlas provides three complementary outputs: tissue expression profiles, a tissue-specificity score (the tau index, ranging from 0 to 1, with values near 1 indicating expression concentrated in one of the sampled tissues), and co-expression neighbors based on Pearson correlations across samples (by default, the top 20 neighbors with |r| ≥ 0.5). Sample-size categories summarize data coverage at query time; they do not constitute a formal measure of statistical confidence. For cucumber, the ChineseLong V3 matrix spans 24,267 genes across 523 samples from 88 projects (Supplemental Table 3).

The Protein module supports structure-guided functional analysis, providing predicted structures and candidate binding sites across the nine species. We used ESMFold (Lin et al., 2023) to predict structures for all nine species, generating 242,316 models (Supplemental Table 3). These models were paired with GPSite binding-site predictions (Yuan et al., 2024), yielding 14.2 million predicted binding-site residue annotations for ligand categories including proteins, peptides, DNA, RNA, ATP, and metal ions. Each protein structure can be viewed and downloaded, together with per-residue confidence scores (pLDDT), allowing users to inspect predicted folds, confidence profiles, and candidate binding sites without running a separate modeling pipeline. Integration with the Gene module enables protein-altering variants to be examined in the context of annotated domains and predicted binding sites. Together, these combined records allow a researcher to move from an observed variant to a structural and functional hypothesis within a single query, facilitating variant prioritization and functional studies across cucurbit crops.

### A curated Cucurbitaceae literature corpus with hybrid retrieval

The Literature module provides access to a curated corpus of cucurbit research papers through hybrid retrieval. We assembled the corpus through collection, quality filtering, indexing, and daily updating (expanded workflow in Supplemental Figure 4). The collection query combines species names with thematic terms, which inevitably retrieves irrelevant records, including clinical papers containing “watermelon” and echinoderm studies containing “sea cucumber”. Automated blacklist filtering, followed by manual review of each record, yielded a curated corpus of 18,305 cucurbit research papers (as of 8 September 2026; Figure 2D; filtering criteria in Methods). A daily pipeline retrieves newly indexed PubMed records for filtering and integration into the live service.

Short gene symbols and allele strings can be difficult to retrieve using semantic similarity alone. The module therefore combines keyword matching and semantic retrieval, with optional cross-encoder reranking (Chen et al., 2024). Keyword and vector search results are merged, and returned passages are linked to source metadata, including a DOI when available (implementation parameters in Methods). We adopted passage-level hybrid retrieval because our gene-specific queries demand precision on short passages.

### CucurbitBench: an expert benchmark for cucurbit functional genomics

To evaluate CucurbitAgent on questions relevant to cucurbit research, we developed CucurbitBench(Supplemental Note 1), an expert-authored benchmark for cucurbit functional genomics. The benchmark assesses eight capabilities: cross-version gene identifier resolution, gene structure and functional annotation, interpretation of natural-variant consequences, tissue expression and co-expression inference, protein structure and binding-site interpretation, literature citation tracing, cross-module evidence integration, and handling of out-of-domain prompts. Cross-module items require evidence from more than one module, while negative controls probe responses to out-of-domain prompts such as a query about “sea cucumber”. The core set contains 50 English items (Q01–Q50, Supplemental Note 1), covering genes (Q01–Q13), expression (Q14–Q25), proteins (Q26–Q37), literature (Q38–Q45), cross-module integration (Q46–Q48), and negative controls (Q49–Q50; capability coverage in Supplemental Table 4). The items span all nine cultivated cucurbit species. Reference answers were exported directly from the platform’s local databases and verified by domain experts; no model-generated content was used.

We compared CucurbitAgent with nine general-purpose LLMs: Claude-Haiku-4.5, DeepSeek-V3, Gemini-2.5-flash, GLM-4.7-flash, GPT-4.1-mini, Hunyuan-HY3, Kimi-K2, Qwen-plus, and Seed-2.0-mini. Three cucurbit experts scored all responses under blinded conditions. Each response was rated on four dimensions—factual accuracy, completeness, relevance, and traceability—using a 0–100 scale, and the ratings were combined into a weighted composite score (weights in Methods). The traceability criterion penalizes fabricated citations and unverifiable claims, addressing a well-documented concern about LLM hallucinations (Farquhar et al., 2024).

### Performance of CucurbitAgent on CucurbitBench

Blinded expert review yielded an overall win rate of 96.0% for CucurbitAgent against the nine general-purpose LLMs, based on weighted composite scores. Across the eight capability axes, CucurbitAgent scored highest on cross-module evidence integration (94.8), natural-variant interpretation (94.2), and gene structure annotation (93.6), exceeding the mean of the nine control models on every axis but one (Figure 3A; per-module win rates in Supplemental Table 5). The exception was the negative-control class: the system declined queries unrelated to cucurbits, whereas general-purpose LLMs provided more informative explanations of the domain boundary (Figure 3A). Per-question composite scores for CucurbitAgent clustered tightly in the upper range, whereas those of the control models were broadly distributed at much lower values, with Gemini-2.5-flash the strongest baseline (Figure 3B). The advantage of CucurbitAgent was most apparent for queries involving assembly-specific identifiers (e.g., CsaV3_3G027830), natural variants requiring sequence and domain context (e.g., the nonsense variant p.E252* in chalcone synthase), and gene symbols with limited literature coverage (e.g., CsCHS). On these benchmark queries, control models produced incorrect or incomplete answers (per-axis score differences in Supplemental Figure 5; side-by-side examples in Supplemental Figure 6), consistent with documented factual errors in LLM responses (Farquhar et al., 2024). All CucurbitBench items and reference answers are provided in Supplemental Note 2 to support further evaluation and comparisons with additional models.

**Figure 3.**
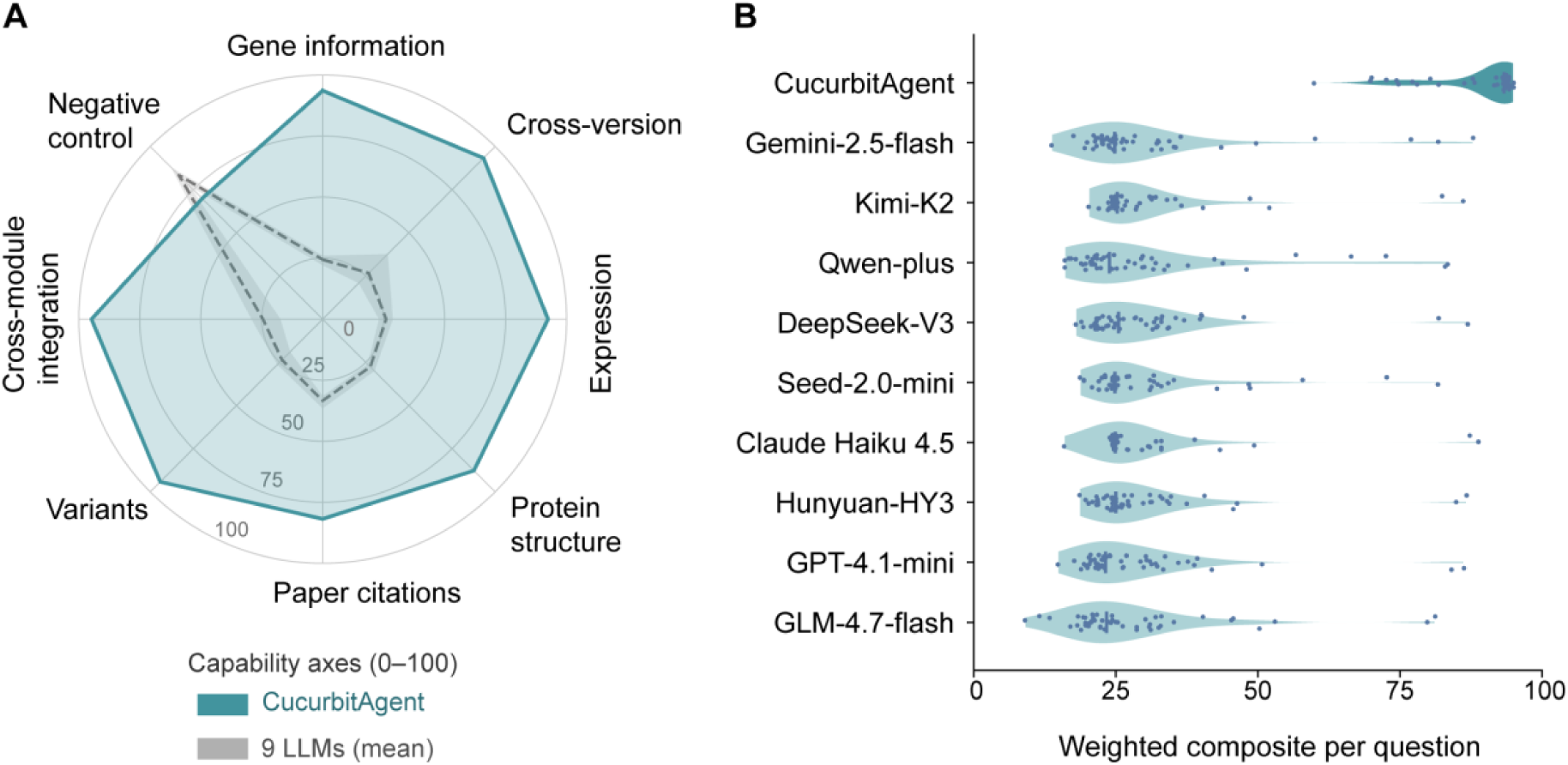
Benchmarking CucurbitAgent against nine general-purpose LLMs on CucurbitBench. (A) Radar chart of scores across eight capability dimensions (0–100), comparing CucurbitAgent (teal) with the mean of the nine general-purpose LLMs (gray dashed line). The gray band indicates the minimum-to-maximum range across control models for each dimension. (B) Distributions of per-question weighted composite scores for the 50 tasks, averaged across three expert raters. Violins show score distributions, and dots represent individual questions.

### An end-to-end case: cucumber chalcone synthase CsCHS

To illustrate how CucurbitAgent operates in practice, we use cucumber chalcone synthase (CsCHS; CsaV3_3G027830) to demonstrate an end-to-end workflow that integrates gene, expression, protein, and literature evidence within a single conversation (Figure 4A; complete session in Supplemental Figure 7). Chalcone synthase catalyzes the first committed step in flavonoid biosynthesis, producing a chalcone precursor for downstream pathways that generate flavones, flavonols, anthocyanins, and proanthocyanidins. Flavonoids contribute to fruit skin and spine pigmentation in cucumber (Li et al., 2020; Xu et al., 2024), connecting specialized metabolism to traits relevant to breeding. CsCHS therefore provides a biologically relevant example for illustrating how the system integrates multiple evidence types for a single gene.

**Figure 4.**
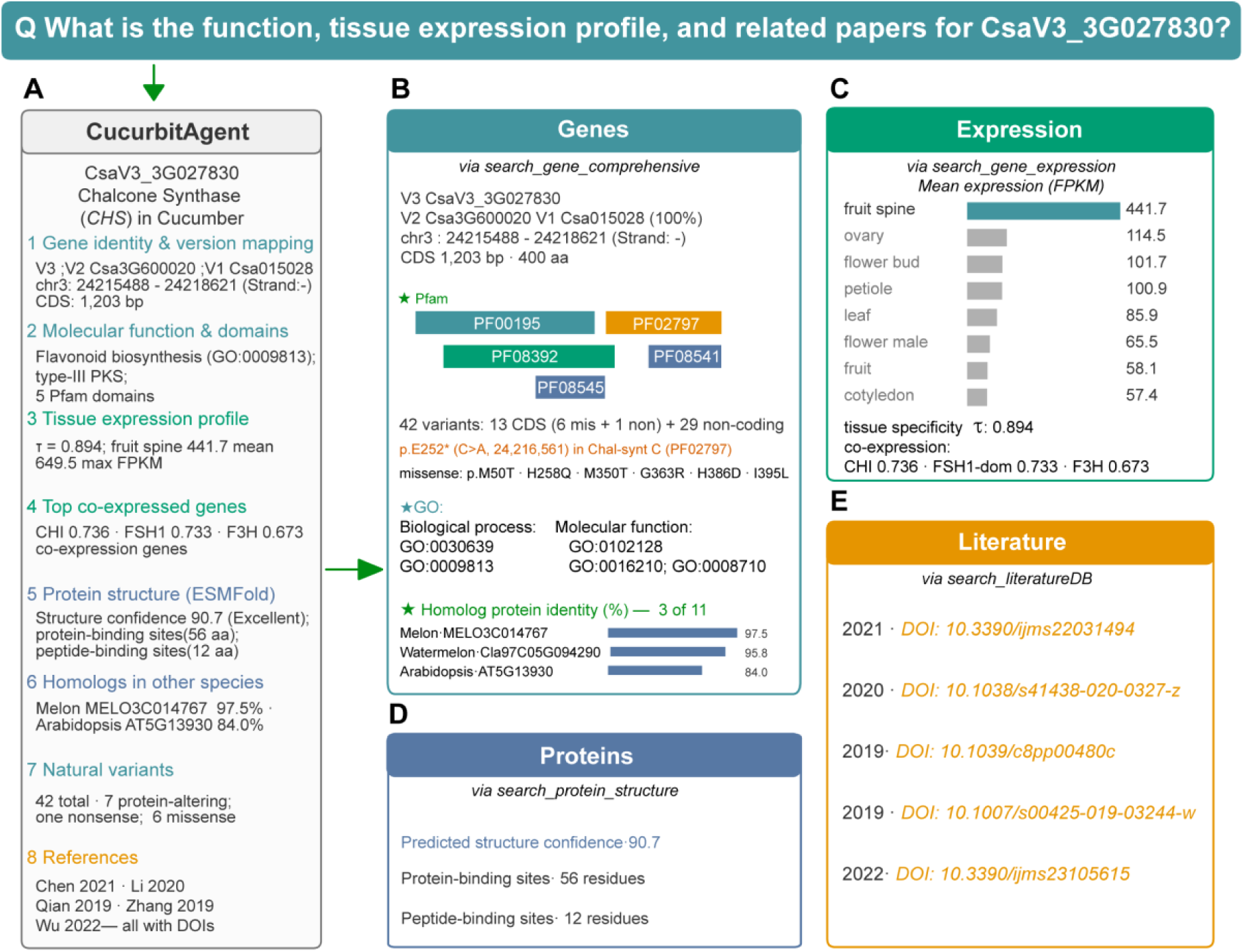
End-to-end case study of cucumber chalcone synthase (CsCHS, CsaV3_3G027830). (A) The agent’s response to “What is the function, tissue expression profile, and related papers for CsaV3_3G027830?” (B) The Gene module resolves the V3/V2/V1 identifiers and reports the genomic location, CDS length, Pfam annotations, and 42 variants. (C) The Expression module shows preferential expression in fruit spines, together with co-expression neighbors from the flavonoid pathway. (D) The Protein module summarizes ESMFold model confidence (pLDDT 90.7) and the numbers of GPSite-predicted protein-and peptide-binding residues. (E) The Literature module returns five related papers with DOIs.

In response to the natural-language prompt (“What is the function, tissue expression profile, and related papers for CsaV3_3G027830?”), the agent retrieves and integrates records from its evidence modules. At the gene level (Figure 4B), CsaV3_3G027830 maps to ChineseLong V2 (Csa3G600020) and V1 (Csa015028), lies at chr3:24,215,488–24,218,621 on the minus strand, and encodes a 400-residue protein with Pfam annotations that include the chalcone/stilbene synthase N-terminal (PF00195) and C-terminal (PF02797) domains. Among the retrieved cucurbit homologs, melon MELO3C014767 and watermelon Cla97C05G094290 share 97.5% and 95.8% sequence identity with CsCHS, respectively; homologs were also retrieved from the other cultivated species and from Arabidopsis (AT5G13930, 84.0%). The natural-variant index reports 42 variants at this locus, including seven predicted to alter the protein sequence: six missense variants and one nonsense variant, p.E252*. In the expression atlas (Figure 4C), CsCHS shows a pronounced tissue bias (tau = 0.894), with the highest mean expression in fruit spines (441.7 FPKM), above that in ovaries, flower buds, and leaves. Its top co-expression neighbors include two flavonoid-pathway enzymes—chalcone-flavanone isomerase (r = 0.736) and flavanone 3-hydroxylase (r = 0.673)—and an FSH1 domain-containing protein (r = 0.733). These associations are consistent with a role for CsCHS in flavonoid biosynthesis in fruit spines. The Protein module returns an ESMFold model (pLDDT 90.7) and GPSite predictions of 56 protein-binding and 12 peptide-binding residues (Figure 4D), allowing p.H258Q to be inspected relative to candidate interaction sites. The p.E252* variant introduces a stop codon at position 252 and is predicted to produce a 251-residue polypeptide, removing residues 252–400 (149 residues), including much of the annotated C-terminal domain PF02797 (residues 239–389). This truncation suggests impaired function, but its effects on transcript abundance, protein stability, and enzymatic activity require experimental validation. The Literature module supplies relevant papers with DOIs (Figure 4E), completing the evidence trail. Within a single conversation, CucurbitAgent brings together evidence otherwise distributed across CuGenDB (Zheng et al., 2019), Phytozome (Goodstein et al., 2012), NCBI, and the primary literature. The CsCHS case illustrates how the CucurbitAgent architecture organizes candidate-gene information and supports the formulation of testable hypotheses.

## DISCUSSION

Large language models are changing how researchers interact with biological knowledge, but their use in specialized crop research remains constrained by factual errors and limited source traceability. Here, we present CucurbitAgent, an evidence-grounded research agent for the Cucurbitaceae. Its unified knowledge base spans nine cultivated species, 255,874 genes, more than 16 million natural variants, 242,316 predicted protein structures with over 14 million predicted binding-site residue annotations, and a curated corpus of over 18,000 publications. Responses are linked to source records for verification, and the platform is freely accessible through an interface supporting 37 languages reaching more than 70% of the global population. To our knowledge, CucurbitAgent is the first research agent dedicated to functional genomics in the Cucurbitaceae.

A central question was whether an agent connected to domain-specific resources could outperform general-purpose LLMs on questions from a specialized crop community. On CucurbitBench, which comprises 50 tasks across eight capability dimensions, CucurbitAgent achieved an overall pairwise win rate of 96.0% against nine general-purpose LLMs based on weighted composite scores from blinded expert assessment. This advantage supports the value of combining curated cucurbit resources with tool-assisted retrieval to answer specialized research questions. Several system features offer plausible explanations for this advantage. Structural records provide context for interpreting protein-altering variants, passage-level citations support verification of literature claims, and cross-version identifier maps connect legacy identifiers to the reference assemblies used by the platform. These features address challenges also motivating broader plant-science agents (Yu et al., 2026).

These results should be interpreted in the context of several limitations. First, grounding answers in curated records reduces unsupported generation, but answer quality still depends on the knowledge base itself, whose completeness, accuracy, and currency require sustained curation. Although daily literature ingestion partially mitigates this issue, the knowledge base must also be updated with newly released genome assemblies and additional transcriptomic datasets to maintain its currency and reliability (Falco et al., 2021). Second, the structural layer relies on computational prediction. ESMFold structures and GPSite binding-site annotations vary in reliability across proteins and residues (Jumper et al., 2021; Lane, 2023; Lin et al., 2023; Yuan et al., 2024), and they support functional hypotheses rather than experimental validation. The modular design allows these resources to be replaced as better predictors emerge, though each replacement will require integration and validation. Third, data coverage remains uneven. Publicly available natural-variant data currently cover only five of the nine species, and the numbers of publicly available RNA-seq samples vary widely among species, with heterogeneity among projects potentially affecting expression and co-expression summaries. In addition, the literature corpus currently consists of English-indexed journals. These gaps define directions for future expansion. Finally, CucurbitBench currently contains 50 English items; larger, independently curated benchmarks will be needed to assess generalizability.

CucurbitAgent offers a unified interface for questions that otherwise require separate searches of CuGenDB, Phytozome, NCBI, and the primary literature. The platform presents candidate-gene information together with its sources, reducing the effort needed to retrieve and organize multi-omics evidence. Its 37-language interface also broadens access for researchers who do not work in English. The architecture itself is the more general contribution. It rests on reusable components rather than on cucurbits: a reconciled multi-omics knowledge base, module-specific retrieval tools, and a domain boundary that keeps every answer verifiable. Other crops can assemble these components into comparable agents. So can any specialized field with curated, queryable records. The framework can also accommodate future language models without redesign. We expect this evidence-grounded paradigm to accelerate gene-function discovery and its translation into breeding across the world’s crop diversity.

## METHODS

### Literature corpus collection, filtering, and daily updates

PubMed and Web of Science were queried using the scientific names of the nine cultivated species (Cucumis sativus, C. melo, Citrullus lanatus, Cucurbita moschata, C. pepo, Momordica charantia, Lagenaria siceraria, Luffa cylindrica, Benincasa hispida), their common names, Cucurbitaceae genus names, and terms related to genetics, breeding, expression, metabolism, and molecular experiments. Retrieved records were filtered in two stages. First, an automated keyword blacklist excluded off-target records, including echinoderm studies (e.g., sea cucumber), studies of other crop families, and records matched through ambiguous names (e.g., “watermelon stomach”). Each remaining record was then manually reviewed for relevance to cucurbit biology. After both stages, the corpus contained 18,305 cucurbit research papers as of September 8, 2026. An automated pipeline applies the same filtering procedure to newly deposited PubMed records each day, keeping the corpus current with the latest research.

### Implementation of literature retrieval

Research queries often contain short gene symbols, assembly-specific identifiers (e.g., CsaV3_3G027830), or allele notation that may be poorly represented by semantic embeddings. The module therefore combines keyword and semantic retrieval in a hybrid strategy. Abstracts and available full texts are divided into chunks using LangChain RecursiveCharacterTextSplitter (chunk_size = 512 characters, overlap = 60 characters). The chunks are indexed in two complementary systems: a SQLite FTS5 full-text index for keyword matching, and a ChromaDB vector index using BAAI/bge-m3 embeddings (Chen et al., 2024) for semantic retrieval. At query time, the top 40 candidates are retrieved from each index and merged by reciprocal rank fusion (RRF, k = 60), which also removes duplicates. An optional cross-encoder (BAAI/bge-reranker-base) then reranks the top 30 fused candidates, and the 10 highest-ranked chunks are returned with paper metadata (title, authors, year, PMID/DOI). Each returned chunk is linked to its source paper and passage, with PMID or DOI metadata where available.

### Gene and variant database construction

Reference genome sequences, gene structures, protein sequences, and functional annotations for the nine cultivated species were obtained from community releases and standardized into species-specific master tables following CuGenDB/NCBI conventions (Goodstein et al., 2012; Howe et al., 2020; Yu et al., 2023; Zheng et al., 2019). The reference assemblies and associated publications were as follows: cucumber ChineseLong v3 (Li et al., 2019; v4 in Guan et al., 2024; v2 in Li et al., 2011; v1 in Huang et al., 2009); melon DHL92 v4 (Castañera et al., 2020; v3.5.1 in Argyris et al., 2015; v3.6.1 in Ruggieri et al., 2018; v3.5 in Garcia-Mas et al., 2012); watermelon 97103 v2.5 (assembly in Guo et al., 2019; v1 in Guo et al., 2013; T2T super-pangenome in Zhang et al., 2024; population pangenome in Sun et al., 2026); pumpkin C. moschata Rifu v1/v2 (Sun et al., 2017) and C. pepo MU-CU-16 v4.1 (Montero-Pau et al., 2018); bitter gourd OHB3-1 v2/Goya (Matsumura et al., 2020; v1 in Urasaki et al., 2017); bottle gourd USVL1VR-Ls v1 (Wu et al., 2017); sponge gourd L. cylindrica (Zhang et al., 2020); and wax gourd B227 (Xie et al., 2019) (Supplemental Table 3). GO and Pfam annotations were added from species-specific GAF files and domain tables (Gene Ontology Consortium, 2021; Mistry et al., 2021).

Cross-version gene identifiers are reconciled using explicit mapping files for cucumber V1–V3, watermelon V1–V2.5, melon V3.5.1–V4, and C. moschata V1–V2. For each species, mappings are anchored to the latest assembly version supported by the platform—for cucumber, V1 and V2 identifiers are aligned to V3 gene loci (Supplemental Table 3)—so that legacy identifiers from older publications resolve to the platform’s reference assembly. For identifiers covered by these maps, the module returns a unified gene card with corresponding identifiers across assemblies, chromosomal coordinates, exon–intron structure, GO and Pfam annotations, natural variants, and candidate homologs. This approach reduces ambiguity when multiple assembly versions coexist.

The natural-variant index is organized by gene locus. For the five species currently covered— cucumber, watermelon, melon, C. pepo, and C. moschata—the index contains 16.0 million variant sites (cucumber maps from Qi et al., 2013 and Li et al., 2022; watermelon from Guo et al., 2019 and Sun et al., 2026; melon GBS from Wang et al., 2021; Cucurbita C. pepo and C. moschata GBS from Yu et al., 2023), with each site recording its alleles and, for protein-altering variants, the resulting amino-acid change. Variants are classified by coding-sequence (CDS) overlap, and coding substitutions are further classified as synonymous or protein-altering (nonsynonymous). The cucumber and watermelon datasets derive from whole-genome resequencing and contain millions of SNPs per species. The melon, C. pepo, and C. moschata datasets derive from genotyping-by-sequencing (GBS), which samples regions near restriction-enzyme cut sites, and contain tens of thousands of SNPs per species. Cross-species candidate homologs were identified using DIAMOND blastp (Buchfink et al., 2021) across the nine cucurbit species and four model plants (Arabidopsis, maize, rice, and tomato). Hits with E-value ≤ 1 × 10^-5^, identity ≥ 40%, and query coverage ≥ 50% were retained in a family-wide sequence-similarity table to support comparative functional analyses.

### Expression and co-expression database construction

Public RNA-seq expression matrices (FPKM) were compiled for the nine cultivated species. Metadata were organized by species, project, and tissue to support tissue-specific queries. The collection contains 1,457 RNA-seq samples across the nine species; the cucumber ChineseLong V3 matrix contains 24,267 genes × 523 samples from 88 projects (Supplemental Table 3).

Data availability is summarized by four sample-size categories: high (≥200 samples), moderate (100–199), limited (50–99), and sparse (<50). These categories indicate sample counts rather than calibrated statistical confidence. Tissue expression profiles summarize mean FPKM by tissue; tissue specificity is quantified using the tau index (Yanai et al., 2005),

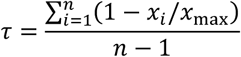

where *x_i_* is the mean FPKM of a gene in tissue *i* and *x*_max_ is its maximum across the *n* profiled tissues; *τ* ranges from 0 (equal expression across the profiled tissues) to 1 (expression restricted to a single profiled tissue). Co-expression neighbors are identified using Pearson correlations between FPKM expression profiles across samples within each species,

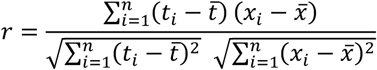

where *t_i_* and *x_i_* are the FPKM values of the target gene and a candidate gene in sample *i*, and *t̄* and *x̄* are their means across the *n* samples. By default, the top 20 neighbors with |r| ≥ 0.5 are returned in interactive tables and bar charts, allowing researchers to identify candidate genes with positively or negatively correlated expression profiles. Correlated expression alone does not establish shared regulation.

### Protein structure and binding-site database construction

Proteome-wide structures for the nine cultivated species were predicted using ESMFold (Lin et al., 2023) and stored in PDB format (Berman et al., 2000; Burley et al., 2025), yielding 242,316 models (Supplemental Table 3). Per-residue confidence scores (pLDDT) are stored in the B-factor field of the PDB files. We then used GPSite (Yuan et al., 2024) to predict binding-site residues in each model. This yielded 14.2 million binding-site residue annotations, covering protein, peptide, DNA, RNA, ATP, and metal-ion ligands. The annotation tables accompany the PDB files and are accessible through both the Protein module and the agent tools. Protein-altering variants retrieved by the Gene module can be mapped to the corresponding residues and examined alongside domain, interface, and predicted binding-site annotations, supporting hypotheses about their functional effects. Together, these structural resources provide a foundation for downstream studies of protein function, ligand interactions, and variant effects across cucurbit crops.

### Tool-calling orchestration of the top-level agent

The top-level session agent uses a ReAct-style tool-calling loop (Yao et al., 2023). At each step, a decision model (default DeepSeek-V3, temperature 0.2) selects one of four module tools: search_gene_comprehensive, search_gene_expression, search_protein_structure, or search_literatureDB. Each tool is exposed through a function-calling interface and returns structured JSON evidence. To limit latency and computational cost, each user turn permits at most seven tool calls (max_tool_calls = 7). Once this limit is reached, the model generates a final synthesis without further tool calls, integrating the retrieved evidence into a natural-language answer with numbered citations. This retrieval-augmented generation approach (Lewis et al., 2020) grounds answers in the built-in knowledge base and reduces fabricated identifiers, coordinates, and citations (Farquhar et al., 2024).

### Direct module queries and domain restrictions

The standalone interfaces of the four modules use a direct retrieve-and-answer workflow. Each module calls its own tool to retrieve structured data and then answers the user’s question in a single step, using the conversational context (temperature 0.3) without invoking the top-level orchestration loop. A domain gatekeeper restricts answers to cucurbit genomics and breeding and refuses out-of-scope queries (e.g., the “sea cucumber” genome) to reduce cross-domain errors. Search and retrieval functions are freely available without limits. Only the AI conversation features require an authenticated session, with daily quotas applied per user (default 50 questions/day) and per IP address (default 1000 questions/day).

### Platform engineering

The CucurbitAgent backend is implemented in Python with FastAPI, and the front end is built with Next.js. The backend handles module retrieval, agent orchestration, and database access, while the front end provides gene cards, expression profiles, a protein structure viewer, and literature evidence panels. The conversation endpoint streams tokens, status updates, and tool-call traces to the browser through server-sent events; a “thinking process” panel displays these retrieval and generation events in real time, so that users can follow the agent’s workflow. Gene, variant, and expression master tables are stored in a relational database, literature embeddings in ChromaDB, the full-text index in SQLite FTS5, and PDB structure files in object storage. Together, these storage backends serve the four evidence modules.

### Multilingual text interface

The portal accepts typed queries in 37 languages and returns answers in the user’s language, while preserving retrieved evidence, gene identifiers, and citations. Query and answer translation are implemented as preprocessing and postprocessing steps around the agent, so that all supported languages share the same evidence-retrieval, tool-calling, and citation-linking workflow.

### CucurbitBench evaluation protocol

CucurbitBench covers eight capability dimensions: cross-version gene identifier resolution, gene structure and functional annotation, natural-variant consequence interpretation, tissue expression and co-expression inference, protein structure and binding-site interpretation, literature citation tracing, cross-module evidence integration, and negative controls that test whether the system refuses out-of-domain prompts rather than fabricating an answer. The core set contains 50 English items (Q01–Q50) and covers all nine cultivated cucurbit species. Category assignments are summarized in Supplemental Note 1 and Supplemental Table 4.

Reference answers were exported from the platform’s local database and verified by domain experts. Three cucurbit specialists, blinded to the response source, independently scored each answer on four dimensions using a 0–100 scale: factual accuracy (40%), completeness (20%), relevance (20%), and traceability (20%). The dimension scores were weighted and summed to obtain a composite score (full protocol in Supplemental Table 6).

CucurbitAgent was evaluated using DeepSeek-V3 with the platform’s standard configuration. The nine general-purpose control models (GPT-4.1-mini, Gemini-2.5-flash, Claude-Haiku-4.5, DeepSeek-V3, Kimi-K2, Seed-2.0-mini, Qwen-plus, GLM-4.7-flash, and Hunyuan-HY3) were accessed through a unified API gateway (OpenRouter) with default decoding settings, and answered the same questions without access to the platform’s local cucurbit databases. This design compares a retrieval-equipped system built on curated data against general-purpose LLMs. The complete item set, reference answers, model responses, and expert scores are provided in Supplemental Notes 1 and 2 to support community comparisons and evaluations with additional models and larger item sets.

### Statistical analysis

Three expert raters independently scored each response for factual accuracy (A), completeness (C), relevance (R), and traceability (T), each on a 0–100 scale. For each benchmark item, scores were averaged across the three raters within each dimension. The composite score was then calculated as a weighted sum:

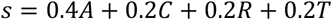

For each item, CucurbitAgent’s composite score was compared with that of each of the nine control models, yielding 50 × 9 = 450 item-level comparisons. The overall win rate (WR) was defined as the proportion of comparisons in which CucurbitAgent’s composite score was strictly higher than that of the control model; ties did not count as wins:

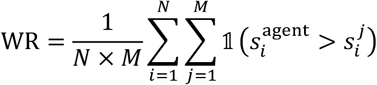

where *N* = 50 is the number of benchmark items, *M* = 9 is the number of control models, s_*i*_^agent^ and s_*i*_^*j*^ denote the composite scores of CucurbitAgent and control model *j* on item *i*, respectively, and 1(⋅) is the indicator function. Confidence intervals were estimated using a nonparametric bootstrap: the 50 items were resampled with replacement 10,000 times, the win rate was recomputed for each resample, and the 95% confidence interval was taken as the 2.5th and 97.5th percentiles of the resulting bootstrap distribution. To verify the reliability of the expert scores and the statistical robustness of the win rate, we performed two complementary analyses. First, inter-rater agreement was high, with Krippendorff’s α of 0.74 for control responses and 0.83 for CucurbitAgent responses, indicating that the expert ratings were consistent and reproducible. Second, item-level composite scores were compared between CucurbitAgent and each control model using two-sided Wilcoxon signed-rank tests, confirming that the advantage of CucurbitAgent was statistically significant against every control model (P < 0.001 in all nine comparisons).

## DATA AND CODE AVAILABILITY

The CucurbitAgent is currently accessible at https://cucurbitagent.com. Source code, the CucurbitBench dataset (items, reference answers, model responses, and expert scores), and knowledge-base snapshots are available at https://github.com/sevencheung2021/CucurbitAgent.

## AUTHOR CONTRIBUTIONS

C.W. conceived the study. R.Z. and C.W. drafted the manuscript. R.Z. led platform engineering and bioinformatics integration. Jian Zhang contributed to module design, data collection and integration, and manuscript revision. Y.Z. contributed to literature curation. J.Y., X.Z., C.X., Jialong Zhang, and H.Z. contributed to discussions and domain validation. All authors reviewed and approved the final manuscript.

## ACKNOWLEDGMENTS

We thank our colleagues at the Beijing Vegetable Research Center, Beijing Academy of Agriculture and Forestry Sciences (BVRC/BAAFS), for discussions on cucurbit breeding applications and feedback on the CucurbitAgent interface. We thank Prof. Shaofang Li (BVRC/BAAFS) for providing computational resources and data-hosting support. We also thank the many researchers who tested the platform and offered constructive suggestions.

## Supplemental information

Supplemental information is available online with this article.

## REFERENCES

Argyris, J.M., Ruiz-Herrera, A., Madriz-Masis, P., Sanseverino, W., Morata, J., Pujol, M., Ramos-Onsins, S.E., and Garcia-Mas, J. (2015). Use of targeted SNP selection for an improved anchoring of the melon (Cucumis melo L.) scaffold genome assembly. BMC Genomics 16, 4. 10.1186/s12864-014-1196-3.

Berman, H.M., Westbrook, J., Feng, Z., Gilliland, G., Bhat, T.N., Weissig, H., Shindyalov, I.N., and Bourne, P.E. (2000). The Protein Data Bank. Nucleic Acids Res. 28, 235–242. 10.1093/nar/28.1.235.

Bolton, E., Venigalla, A., Yasunaga, M., Hall, D., Xiong, B., Lee, T., Daneshjou, R., Frankle, J., Liang, P., Carbin, M., and Manning, C.D. (2024). BioMedLM: A 2.7B Parameter Language Model Trained On Biomedical Text. arXiv. 10.48550/arxiv.2403.18421.

Buchfink, B., Reuter, K., and Drost, H.-G. (2021). Sensitive protein alignments at tree-of-life scale using DIAMOND. Nat. Methods 18, 366–368. 10.1038/s41592-021-01101-x.

Burley, S.K., Bhatt, R., Bhikadiya, C., Bi, C., Biester, A., Biswas, P., Bittrich, S., Blaumann, S., Brown, R., Chao, H., et al. (2025). Updated resources for exploring experimentally-determined PDB structures and Computed Structure Models at the RCSB Protein Data Bank. Nucleic Acids Res. 53, D564–D574. 10.1093/nar/gkae1091.

Castañera, R., Ruggieri, V., Pujol, M., Garcia-Mas, J., and Casacuberta, J.M. (2020). An improved melon reference genome with single-molecule sequencing uncovers a recent burst of transposable elements with potential impact on genes. Front. Plant Sci. 10, 1815. 10.3389/fpls.2019.01815.

Chen, J., Xiao, S., Zhang, P., Luo, K., Lian, D., and Liu, Z. (2024). M3-Embedding: Multi-Linguality, Multi-Functionality, Multi-Granularity Text Embeddings Through Self-Knowledge Distillation. Findings of the Association for Computational Linguistics: ACL 2024, 2318–2335. 10.18653/v1/2024.findings-acl.137.

Depuydt, T., De Rybel, B., and Vandepoele, K. (2023). Charting plant gene functions in the multi-omics and single-cell era. Trends Plant Sci. 28, 283–296. 10.1016/j.tplants.2022.09.008.

Eberhard, D.M., Simons, G.F., and Fennig, C.D. (eds.). (2025). Ethnologue: Languages of the World, 28th ed. Dallas, TX: SIL International. https://www.ethnologue.com.

Edge, D., Trinh, H., Cheng, N., Bradley, J., Chao, A., Mody, A., Truitt, S., Metropolitansky, D., Ness, R.O., and Larson, J. (2024). From local to global: A graph RAG approach to query-focused summarization. Preprint at arXiv. 10.48550/arXiv.2404.16130.

Falco, G., Shneiderman, B., Badger, J., Carrier, R., Dahbura, A., Danks, D., Eling, M., Goodloe, A., Gupta, J., Hart, C., et al. (2021). Governing AI safety through independent audits. Nat. Mach. Intell. 3, 566–571. 10.1038/s42256-021-00370-7.

Farquhar, S., Kossen, J., Kuhn, L., and Gal, Y. (2024). Detecting hallucinations in large language models using semantic entropy. Nature 630, 625–630. 10.1038/s41586-024-07421-0.

Garcia-Mas, J., Benjak, A., Sanseverino, W., Bourgeois, M., Mir, G., González, V.M., Hénaff, E., Câmara, F., Cozzuto, L., Lowy, E., et al. (2012). The genome of melon (Cucumis melo L.). Proc. Natl. Acad. Sci. USA 109, 11872–11877. 10.1073/pnas.1205415109.

Gene Ontology Consortium. (2021). The Gene Ontology resource: enriching a GOld mine. Nucleic Acids Res. 49, D325–D334. 10.1093/nar/gkaa1113.

Goodstein, D.M., Shu, S., Howson, R., Neupane, R., Hayes, R.D., Fazo, J., Mitros, T., Dirks, W., Hellsten, U., Putnam, N., et al. (2012). Phytozome: a comparative platform for green plant genomics. Nucleic Acids Res. 40, D1178–D1186. 10.1093/nar/gkr944.

Grumet, R., McCreight, J.D., McGregor, C., Weng, Y., Mazourek, M., Reitsma, K., Labate, J., Davis, A., and Fei, Z. (2021). Genetic Resources and Vulnerabilities of Major Cucurbit Crops. Genes 12, 1222. 10.3390/genes12081222.

Guan, J., Miao, H., Zhang, Z., Dong, S., Zhou, Q., Liu, X., Beckles, D.M., Gu, X., Huang, S., and Zhang, S. (2024). A near-complete cucumber reference genome assembly and Cucumber-DB, a multi-omics database. Mol. Plant 17, 1178–1182. 10.1016/j.molp.2024.06.012.

Guo, S., Zhang, J., Sun, H., Salse, J., Lucas, W.J., Zhang, H., Zheng, Y., Mao, L., Ren, Y., Wang, Z., et al. (2013). The draft genome of watermelon (Citrullus lanatus) and resequencing of 20 diverse accessions. Nat. Genet. 45, 51–58. 10.1038/ng.2470.

Guo, S., Zhao, S., Sun, H., Wang, X., Wu, S., Lin, T., Ren, Y., Gao, L., Deng, Y., Zhang, J., et al. (2019). Resequencing of 414 cultivated and wild watermelon accessions identifies selection for fruit quality traits. Nat. Genet. 51, 1616–1623. 10.1038/s41588-019-0518-4.

Howe, K.L., Contreras-Moreira, B., De Silva, N., Maslen, G., Akanni, W., Allen, J., Alvarez-Jarreta, J., Barba, M., Bolser, D.M., Cambell, L., et al. (2020). Ensembl Genomes 2020—enabling non-vertebrate genomic research. Nucleic Acids Res. 48, D689–D695. 10.1093/nar/gkz890.

Huang, S., Li, R., Zhang, Z., Li, L., Gu, X., Fan, W., Lucas, W.J., Wang, X., Xie, B., Ni, P., et al. (2009). The genome of the cucumber, Cucumis sativus L. Nat. Genet. 41, 1275–1281. 10.1038/ng.475.

Jumper, J., Evans, R., Pritzel, A., Green, T., Figurnov, M., Ronneberger, O., Tunyasuvunakool, K., Bates, R., Žídek, A., Potapenko, A., et al. (2021). Highly accurate protein structure prediction with AlphaFold. Nature 596, 583–589. 10.1038/s41586-021-03819-2.

Lane, T.J. (2023). Protein structure prediction has reached the single-structure frontier. Nat. Methods 20, 170–173. 10.1038/s41592-022-01760-4.

Lewis, P., Perez, E., Piktus, A., Petroni, F., Karpukhin, V., Goyal, N., Küttler, H., Lewis, M., Yih, W.-T., Rocktäschel, T., et al. (2020). Retrieval-augmented generation for knowledge-intensive NLP tasks. Advances in Neural Information Processing Systems 33, 9459–9474. https://proceedings.neurips.cc/paper/2020/hash/6b493230205f780e1bc26945df7481e5-Abstract.html.

Li, H., Wang, S., Chai, S., Yang, Z., Zhang, Q., Xin, H., Xu, Y., Lin, S., Chen, X., Yao, Z., et al. (2022). Graph-based pan-genome reveals structural and sequence variations related to agronomic traits and domestication in cucumber. Nat. Commun. 13, 682. 10.1038/s41467-022-28362-0.

Li, J., Luan, Q., Han, J., Zhang, C., Liu, M., and Ren, Z. (2020). CsMYB60 directly and indirectly activates structural genes to promote the biosynthesis of flavonols and proanthocyanidins in cucumber. Hortic. Res. 7, 103. 10.1038/s41438-020-0327-z.

Li, J., Jia, Y., Li, F., Su, X., Luo, J., Dong, Y., Sun, C., Cui, Q., Wang, L., Li, A., et al. (2026). An AI-powered knowledge hub for potato functional genomics. Plant Commun. 7, 101730. 10.1016/j.xplc.2026.101730.

Li, Q., Li, H., Huang, W., Xu, Y., Zhou, Q., Wang, S., Ruan, J., Huang, S., and Zhang, Z. (2019). A chromosome-scale genome assembly of cucumber (Cucumis sativus L.). GigaScience 8, giz072. 10.1093/gigascience/giz072.

Li, Z., Zhang, Z., Yan, P., Huang, S., Fei, Z., and Lin, K. (2011). RNA-Seq improves annotation of protein-coding genes in the cucumber genome. BMC Genomics 12, 540. 10.1186/1471-2164-12-540.

Lin, Z., Akin, H., Rao, R., Hie, B., Zhu, Z., Lu, W., Smetanin, N., Verkuil, R., Kabeli, O., Shmueli, Y., et al. (2023). Evolutionary-scale prediction of atomic-level protein structure with a language model. Science 379, 1123–1130. 10.1126/science.ade2574.

Luo, R., Sun, L., Xia, Y., Qin, T., Zhang, S., Poon, H., and Liu, T.-Y. (2022). BioGPT: generative pre-trained transformer for biomedical text generation and mining. Brief. Bioinform. 23, bbac409. 10.1093/bib/bbac409.

Marks, R.A., Hotaling, S., Frandsen, P.B., and VanBuren, R. (2021). Representation and participation across 20 years of plant genome sequencing. Nat. Plants 7, 1571–1578. 10.1038/s41477-021-01031-8.

Matsumura, H., Hsiao, M.-C., Lin, Y.-P., Toyoda, A., Taniai, N., Tarora, K., Urasaki, N., Anand, S.S., Dhillon, N.P.S., Schafleitner, R., and Lee, C.-R. (2020). Long-read bitter gourd (Momordica charantia) genome and the genomic architecture of nonclassic domestication. Proc. Natl. Acad. Sci. USA 117, 14543–14551. 10.1073/pnas.1921016117.

Mistry, J., Chuguransky, S., Williams, L., Qureshi, M., Salazar, G.A., Sonnhammer, E.L.L., Tosatto, S.C.E., Paladin, L., Raj, S., Richardson, L.J., Finn, R.D., and Bateman, A. (2021). Pfam: The protein families database in 2021. Nucleic Acids Res. 49, D412–D419. 10.1093/nar/gkaa913.

Montero-Pau, J., Blanca, J., Bombarely, A., Ziarsolo, P., Esteras, C., Martí-Gómez, C., Ferriol, M., Gómez, P., Jamilena, M., Mueller, L., Picó, B., and Cañizares, J. (2018). De novo assembly of the zucchini genome reveals a whole-genome duplication associated with the origin of the Cucurbita genus. Plant Biotechnol. J. 16, 1161–1171. 10.1111/pbi.12860.

Provart, N.J., Brady, S.M., Parry, G., Schmitz, R.J., Queitsch, C., Bonetta, D., Waese, J., Schneeberger, K., and Loraine, A.E. (2021). Anno genominis XX: 20 years of Arabidopsis genomics. Plant Cell 33, 832–845. 10.1093/plcell/koaa038.

Qi, J., Liu, X., Shen, D., Miao, H., Xie, B., Li, X., Zeng, P., Wang, S., Shang, Y., Gu, X., Du, Y., Li, Y., Lin, T., Yuan, J., Yang, X., Chen, J., Chen, H., Xiong, X., Huang, K., Fei, Z., Mao, L., Tian, L., Städler, T., Renner, S.S., Kamoun, S., Lucas, W.J., Zhang, Z., and Huang, S. (2013). A genomic variation map provides insights into the genetic basis of cucumber domestication and diversity. Nat. Genet. 45, 1510–1515. 10.1038/ng.2801.

Ruggieri, V., Alexiou, K.G., Morata, J., Argyris, J., Pujol, M., Yano, R., Nonaka, S., Ezura, H., Latrasse, D., Boualem, A., et al. (2018). An improved assembly and annotation of the melon (Cucumis melo L.) reference genome. Sci. Rep. 8, 8088. 10.1038/s41598-018-26416-2.

Stark, R., Grzelak, M., and Hadfield, J. (2019). RNA sequencing: the teenage years. Nat. Rev. Genet. 20, 631–656. 10.1038/s41576-019-0150-2.

Sun, H., Wu, S., Zhang, G., Jiao, C., Guo, S., Ren, Y., Zhang, J., Zhang, H., Gong, G., Jia, Z., et al. (2017). Karyotype Stability and Unbiased Fractionation in the Paleo-Allotetraploid Cucurbita Genomes. Mol. Plant 10, 1293–1306. 10.1016/j.molp.2017.09.003.

Sun, H., Zhang, J., Liao, S., Guo, S., Zhou, Z., Zhao, X., Wu, S., Zhao, J., Gong, G., Wang, J., et al. (2026). Population-level super-pangenome reveals genome evolution and empowers precision breeding in watermelon. Nat. Genet. 58, 1176–1185. 10.1038/s41588-026-02598-8.

Sun, Y., Shang, L., Zhu, Q.-H., Fan, L., and Guo, L. (2022). Twenty years of plant genome sequencing: achievements and challenges. Trends Plant Sci. 27, 391–401. 10.1016/j.tplants.2021.10.006.

Tarazona, S., Arzalluz-Luque, Á., and Conesa, A. (2021). Undisclosed, unmet and neglected challenges in multi-omics studies. Nat. Comput. Sci. 1, 395–402. 10.1038/s43588-021-00086-z.

Urasaki, N., Takagi, H., Natsume, S., Uemura, A., Taniai, N., Miyagi, N., Fukushima, M., Suzuki, S., Tarora, K., Tamaki, M., Sakamoto, M., Terauchi, R., and Matsumura, H. (2017). Draft genome sequence of bitter gourd (Momordica charantia), a vegetable and medicinal plant in tropical and subtropical regions. DNA Res. 24, 51–58. 10.1093/dnares/dsw047.

Wang, X., Ando, K., Wu, S., Reddy, U.K., Tamang, P., Bao, K., Hammar, S.A., Grumet, R., McCreight, J.D., and Fei, Z. (2021). Genetic characterization of melon accessions in the U.S. National Plant Germplasm System and construction of a melon core collection. Mol. Hortic. 1, 11. 10.1186/s43897-021-00014-9.

Zhang, T., Ren, X., Zhang, Z., Ming, Y., Yang, Z., Hu, J., Li, S., Wang, Y., Sun, S., Sun, K., Piao, F., and Sun, Z. (2020). Long-read sequencing and de novo assembly of the Luffa cylindrica (L.) Roem. genome. Mol. Ecol. Resour. 20, 511–519. 10.1111/1755-0998.13129

Wu, S., Shamimuzzaman, M., Sun, H., Salse, J., Sui, X., Wilder, A., Wu, Z., Levi, A., Xu, Y., Ling, K.-S., and Fei, Z. (2017). The bottle gourd genome provides insights into Cucurbitaceae evolution and facilitates mapping of a Papaya ring-spot virus resistance locus. Plant J. 92, 963–975. 10.1111/tpj.13722.

Xie, D., Xu, Y., Wang, J., Liu, W., Zhou, Q., Luo, S., Huang, W., He, X., Li, Q., Peng, Q., Yang, X., Yuan, J., Yu, J., Wang, X., Lucas, W.J., Huang, S., Jiang, B., and Zhang, Z. (2019). The wax gourd genomes offer insights into the genetic diversity and ancestral cucurbit karyotype. Nat. Commun. 10, 5158. 10.1038/s41467-019-13185-3.

Xu, X., Zhu, Y., Yuan, Y., Sohail, H., He, S., Ye, Y., Wang, M., Lv, M., Qi, X., Yang, X., et al. (2024). R2R3-MYB transcription factor CsMYB60 controls mature fruit skin color by regulating flavonoid accumulation in cucumber. Plant J. 119, 796–813. 10.1111/tpj.16797.

Yanai, I., Benjamin, H., Shmoish, M., Chalifa-Caspi, V., Shklar, M., Ophir, R., Bar-Even, A., Horn-Saban, S., Safran, M., Domany, E., et al. (2005). Genome-wide midrange transcription profiles reveal expression level relationships in human tissue specification. Bioinformatics 21, 650–659. 10.1093/bioinformatics/bti042.

Yang, F., Kong, H., Ying, J., Chen, Z., Luo, T., Jiang, W., Yuan, Z., Wang, Z., Ma, Z., Wang, S., et al. (2025). SeedLLM·Rice: A large language model integrated with rice biological knowledge graph. Mol. Plant 18, 1118–1129. 10.1016/j.molp.2025.05.013.

Yao, S., Zhao, J., Yu, D., Du, N., Shafran, I., Narasimhan, K., and Cao, Y. (2023). ReAct: Synergizing reasoning and acting in language models. International Conference on Learning Representations. https://openreview.net/forum?id=WE_vluYUL-X.

Yu, H., Zhou, S., Huang, M., Ding, L., Chen, Y., Wang, Y., Ren, Y., Cheng, N., Wang, X., Liang, J., et al. (2026). PlantScience.ai: An LLM-powered virtual scientist for plant science. Mol. Plant 19, 1117–1123. 10.1016/j.molp.2026.03.010.

Yu, J., Wu, S., Sun, H., Wang, X., Tang, X., Guo, S., Zhang, Z., Huang, S., Xu, Y., Weng, Y., et al. (2023). CuGenDBv2: an updated database for cucurbit genomics. Nucleic Acids Res. 51, D1457–D1464. 10.1093/nar/gkac921.

Yuan, Q., Tian, C., and Yang, Y. (2024). Genome-scale annotation of protein binding sites via language model and geometric deep learning. eLife 13, RP93695. 10.7554/eLife.93695.

Zhang, R., Wang, Y., Yang, W., Wen, J., Liu, W., Zhi, S., Li, G., Chai, N., Huang, J., Xie, Y., et al. (2025). PlantGPT: An Arabidopsis-Based Intelligent Agent that Answers Questions about Plant Functional Genomics. Adv. Sci. 12, e03926. 10.1002/advs.202503926.

Zhang, Y., Zhao, M., Tan, J., Huang, M., Chu, X., Li, Y., Han, X., Fang, T., Tian, Y., Jarret, R.L., Lu, D., Chen, Y., Xue, L., Li, X., Qin, G., Li, B., Sun, Y., Deng, X.W., Deng, Y., Zhang, X., and He, H. (2024). Telomere-to-telomere Citrullus super-pangenome provides direction for watermelon breeding. Nat. Genet. 56, 1750–1761. 10.1038/s41588-024-01823-6.

Zheng, Y., Wu, S., Bai, Y., Sun, H., Jiao, C., Guo, S., Zhao, K., Blanca, J., Zhang, Z., Huang, S., et al. (2019). Cucurbit Genomics Database (CuGenDB): a central portal for comparative and functional genomics of cucurbit crops. Nucleic Acids Res. 47, D1128–D1136. 10.1093/nar/gky944.

