## Supplemental Information for "CucurbitAgent: An AI agent for exploring cucurbit multi-omics resources"

### Contents

**Supplemental Figure 1.** Direct search interface of the Gene module, shown for the example query CsaV3_1G000080 (cucumber) without agent involvement.

**Supplemental Figure 2.** Direct search interface of the Expression module (example query: CsaV3_1G000080).

**Supplemental Figure 3.** Direct search interface of the Protein module (example query: CsaV3_1G000080).

**Supplemental Figure 4.** Construction, daily updating, and hybrid retrieval of the literature corpus.

**Supplemental Figure 5.** CucurbitAgent versus nine general-purpose LLMs: overall scores and per-axis comparison on CucurbitBench.

**Supplemental Figure 6.** Representative CucurbitBench responses of CucurbitAgent and general-purpose LLMs.

**Supplemental Figure 7.** Full agent session for the question "What is the function, tissue expression profile, and related papers for CsaV3_3G027830?".

**Supplemental Table 1.** Module-level data inventory of the CucurbitAgent knowledge foundation.

**Supplemental Table 2.** Supported languages of the multilingual text interface.

**Supplemental Table 3.** Data inventory per species across nine cucurbit assemblies.

**Supplemental Table 4.** CucurbitBench capability coverage matrix (50 items).

**Supplemental Table 5.** Per-module win rates of CucurbitAgent against general-purpose LLMs.

**Supplemental Table 6.** Evaluation protocol of CucurbitBench and blinded expert review.

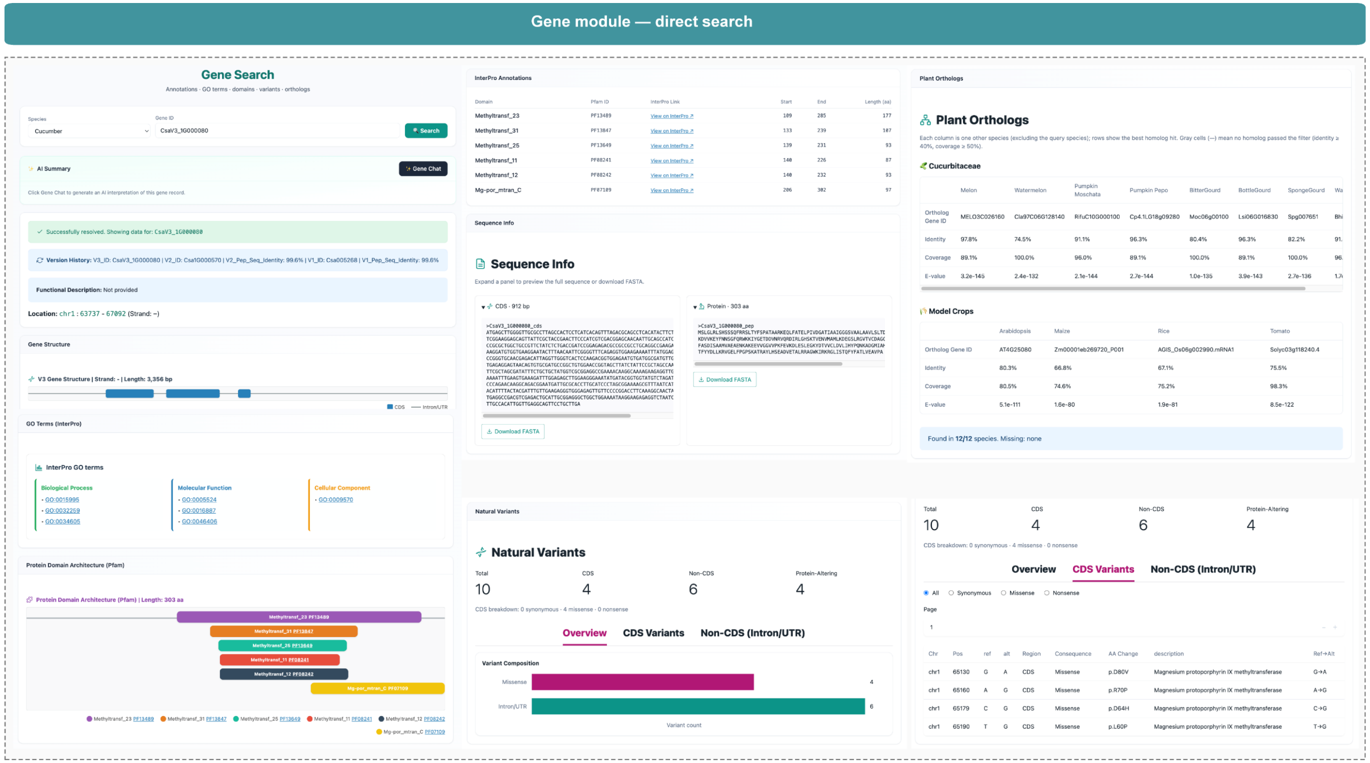


**Supplemental Figure 1. Direct search interface of the Gene module, shown for the example query CsaV3_1G000080 (cucumber) without agent involvement.**

The interface returns the gene card (locus, gene structure, CDS length), cross-version identifier mapping (V3/V2/V1), GO terms and Pfam domain architecture, natural-variant statistics, and best-hit orthologs across the eight other cucurbits and four model species.

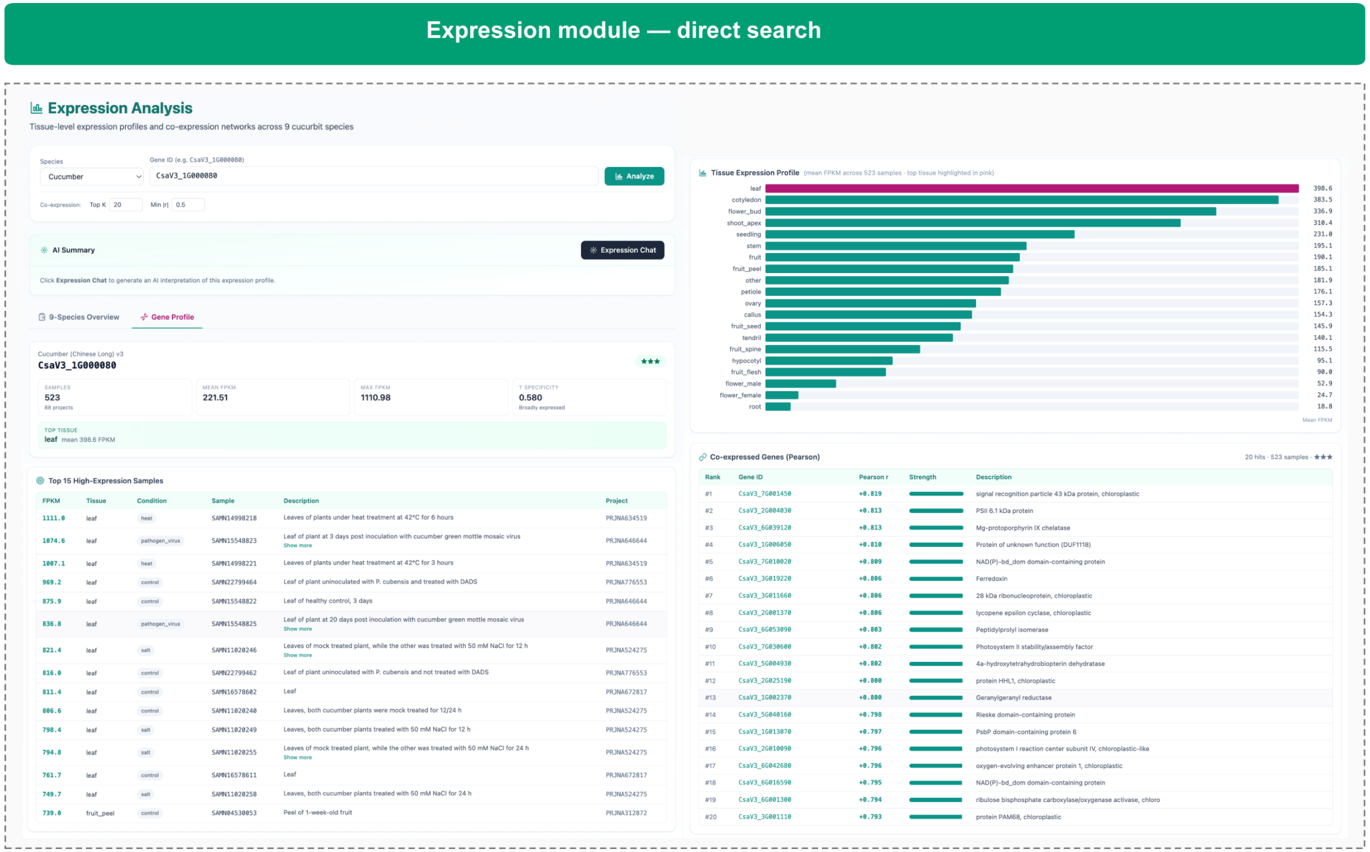


**Supplemental Figure 2. Direct search interface of the Expression module (example query: CsaV3_1G000080).**

The interface provides the gene's expression summary, tissue-level expression profile, top high-expression samples with experimental conditions, and top co-expressed genes ranked by Pearson *r*.

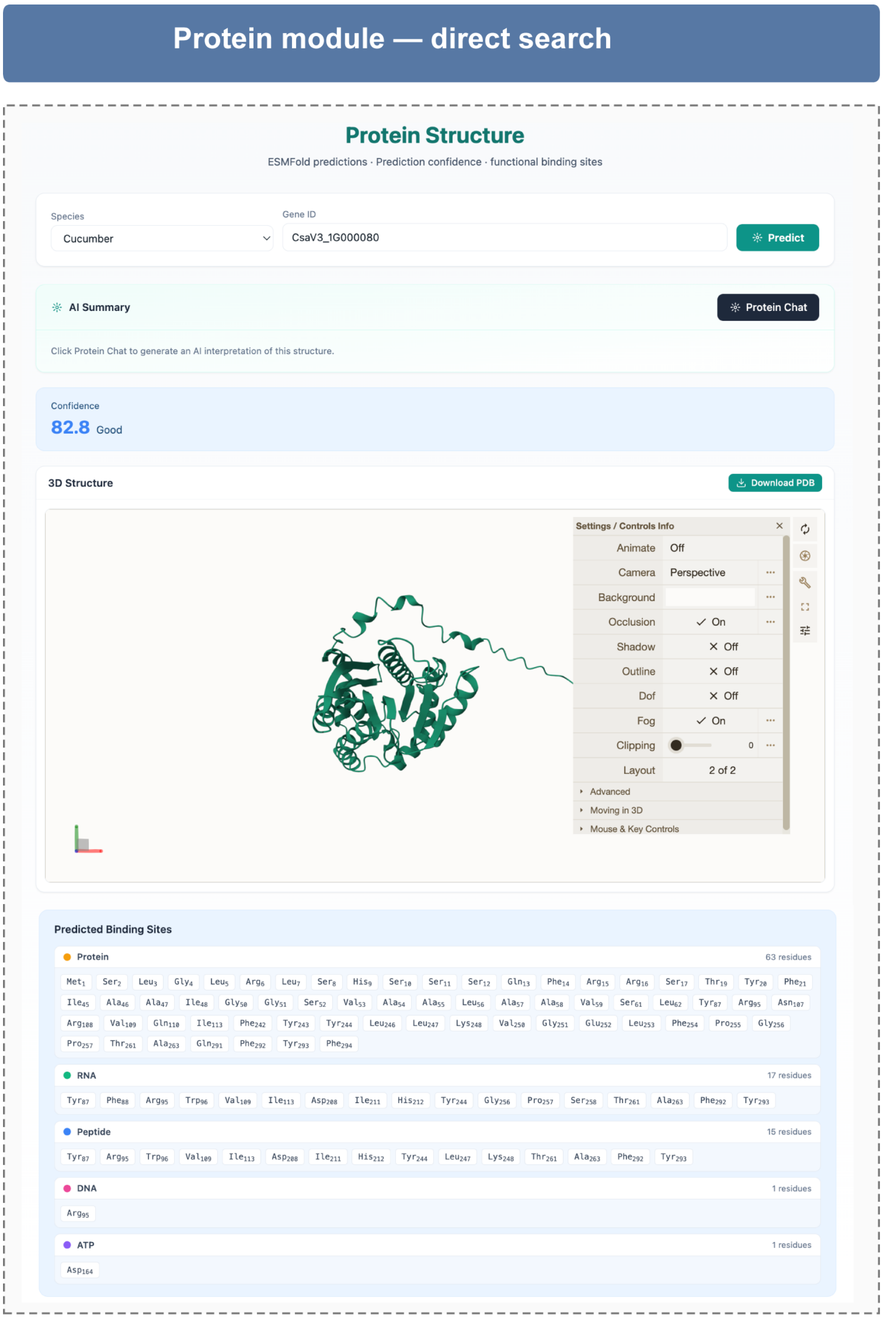


**Supplemental Figure 3. Direct search interface of the Protein module (example query: CsaV3_1G000080).**

The interface returns the ESMFold-predicted structure with a model-confidence score and quality label, an interactive 3D viewer with PDB download, and GPSite-predicted binding sites grouped by ligand type.

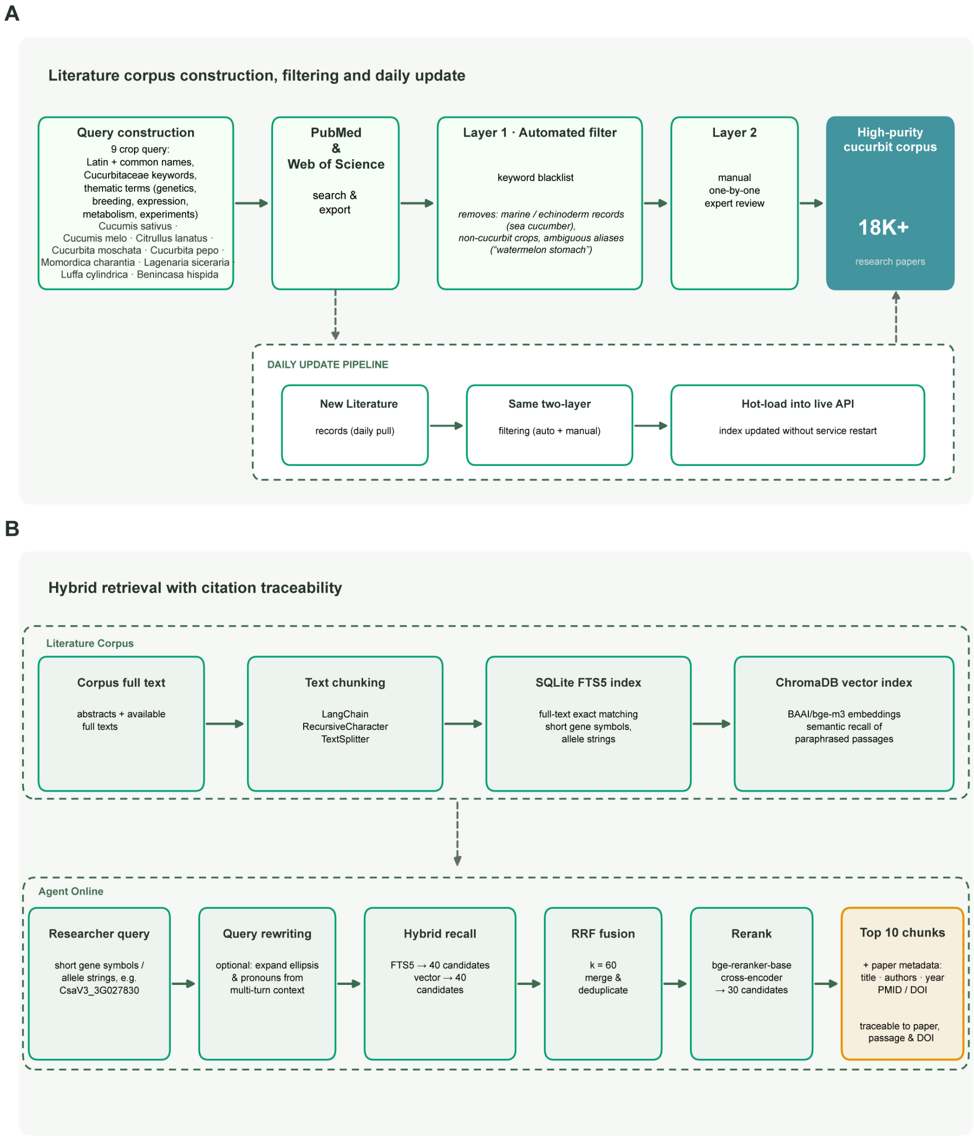


**Supplemental Figure 4. Construction, daily updating, and hybrid retrieval of the literature corpus.**

(A) Corpus construction, filtering, and daily update. PubMed and Web of Science were searched with queries combining the Latin and common names of the nine cultivated cucurbits with Cucurbitaceae keywords and thematic terms. Records are filtered in two layers—an automated keyword blacklist that removes off-topic records (for example, papers on sea cucumbers or on "watermelon stomach", a gastric disease), followed by one-by-one manual expert review—yielding a corpus of 18,305 papers. New records are pulled daily, passed through the same two-layer filter, and hot-loaded into the live API without restarting the service (dashed box).

(B) Hybrid retrieval with citation traceability. Abstracts and available full texts are split into chunks and indexed in parallel in two complementary systems: SQLite FTS5 for exact matching (short gene symbols and allele strings) and ChromaDB for semantic matching of paraphrased passages. Candidates from both indexes are fused, re-ranked, and returned with paper metadata (title, authors, year, PMID/DOI), so that every passage can be traced to its source publication.

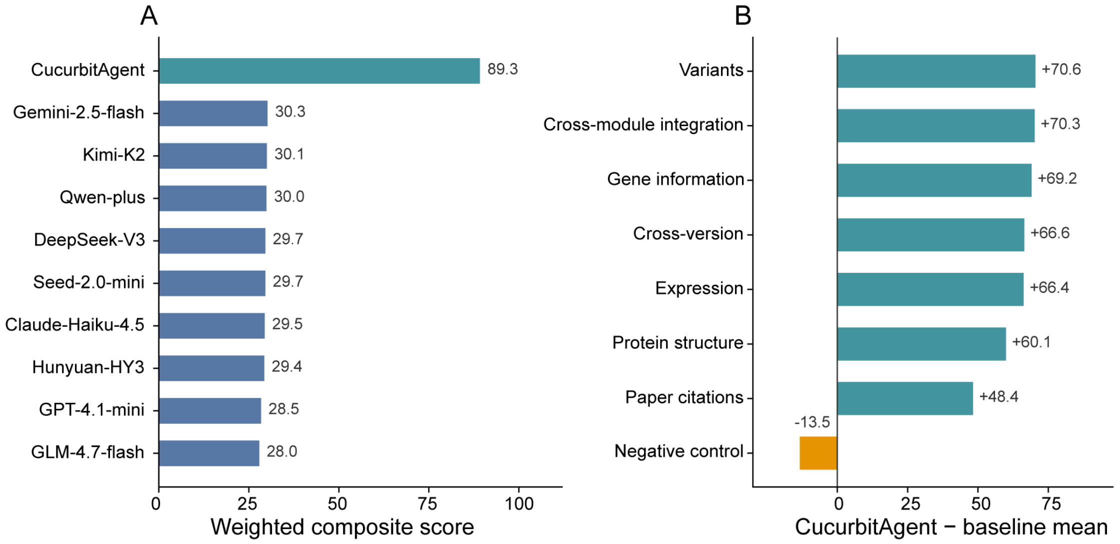


**Supplemental Figure 5. CucurbitAgent versus nine general-purpose LLMs: overall scores and per-axis comparison on CucurbitBench.**

(A) Weighted composite scores of CucurbitAgent and nine general-purpose LLMs (0–100 scale).

(B) Difference between CucurbitAgent and the baseline mean for each capability axis.

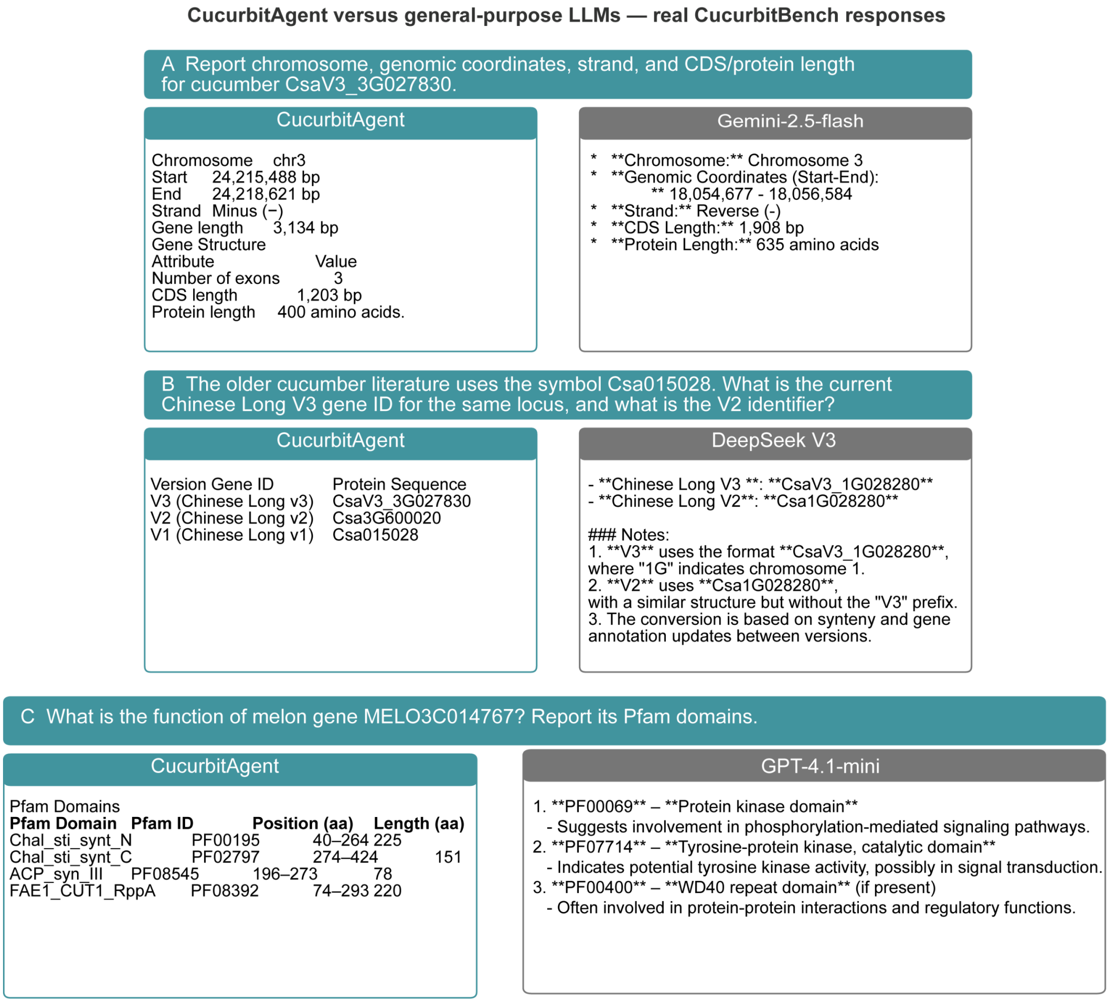


**Supplemental Figure 6. Representative CucurbitBench responses of CucurbitAgent and general-purpose LLMs.**

(A) CucurbitAgent and Gemini-2.5-flash on a gene structure query (cucumber CsaV3_3G027830).

(B) CucurbitAgent and DeepSeek V3 on a gene identifier conversion query (cucumber Csa015028).

(C) CucurbitAgent and GPT-4.1-mini on a protein function and Pfam domain query (melon MELO3C014767).

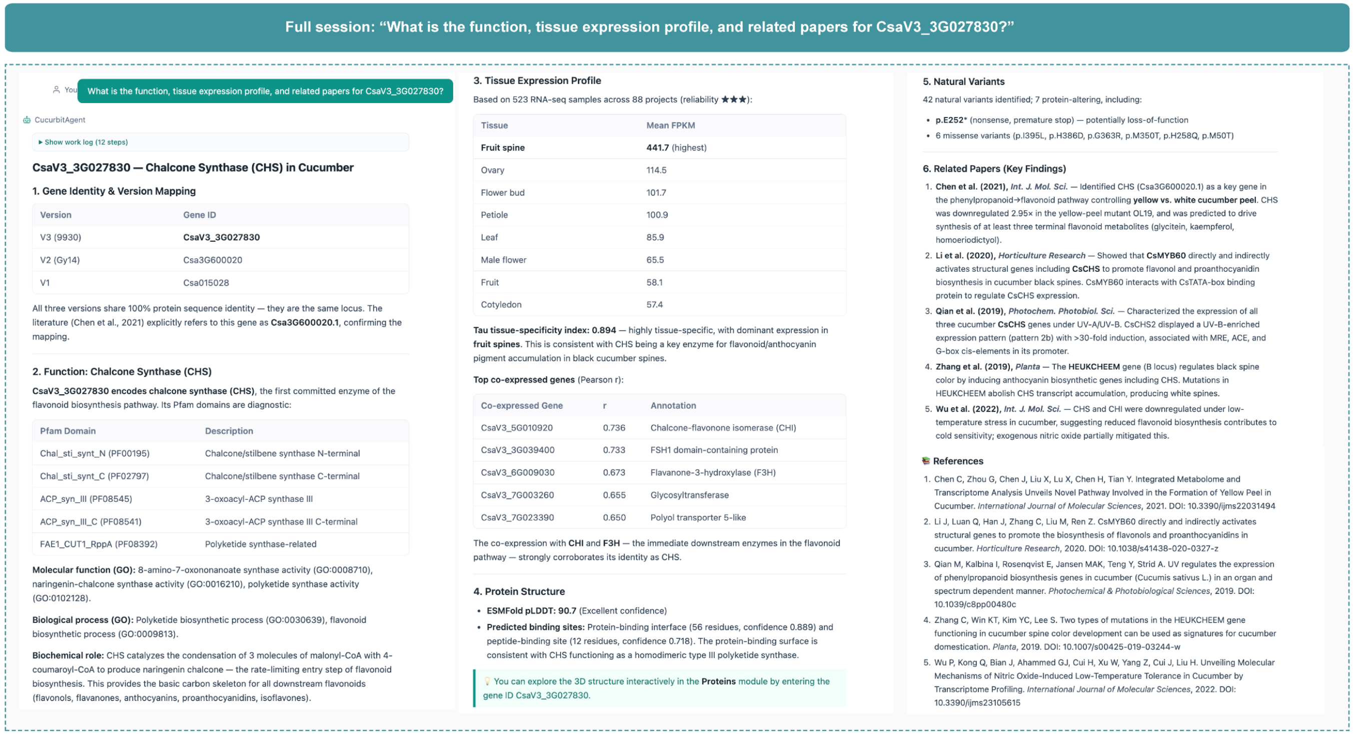

Full session: “What is the function, tissue expression profile, and related papers for CsaV3_3G027830?”

You: What is the function, tissue expression profile, and related papers for CsaV3_3G027830?

CucurbitAgent

▸ Show work log (12 steps)

**CsaV3_3G027830 — Chalcone Synthase (CHS) in Cucumber**

**1. Gene Identity & Version Mapping**

| Version | Gene ID |
|---|---|
| V3 (9930) | **CsaV3_3G027830** |
| V2 (Gy14) | Csa3G600020 |
| V1 | Csa015028 |

All three versions share 100% protein sequence identity — they are the same locus. The literature (Chen et al., 2021) explicitly refers to this gene as **Csa3G600020.1**, confirming the mapping.

**2. Function: Chalcone Synthase (CHS)**

**CsaV3_3G027830 encodes chalcone synthase (CHS)**, the first committed enzyme of the flavonoid biosynthesis pathway. Its Pfam domains are diagnostic:

| Pfam Domain | Description |
|---|---|
| Chal_sti_synt_N (PF00195) | Chalcone/stilbene synthase N-terminal |
| Chal_sti_synt_C (PF02797) | Chalcone/stilbene synthase C-terminal |
| ACP_syn_III (PF08545) | 3-oxoacyl-ACP synthase III |
| ACP_syn_III_C (PF08541) | 3-oxoacyl-ACP synthase III C-terminal |
| FAE1_CUT1_RppA (PF08392) | Polyketide synthase-related |

**Molecular function (GO):** 8-amino-7-oxononanoate synthase activity (GO:0008710), naringenin-chalcone synthase activity (GO:0016210), polyketide synthase activity (GO:0102128).

**Biological process (GO):** Polyketide biosynthetic process (GO:0030639), flavonoid biosynthetic process (GO:0009813).

**Biochemical role:** CHS catalyzes the condensation of 3 molecules of malonyl-CoA with 4-coumaroyl-CoA to produce naringenin chalcone — the rate-limiting entry step of flavonoid biosynthesis. This provides the basic carbon skeleton for all downstream flavonoids (flavonols, flavanones, anthocyanins, proanthocyanidins, isoflavones).

**3. Tissue Expression Profile**

Based on 523 RNA-seq samples across 88 projects (reliability ★★★):

| Tissue | Mean FPKM |
|---|---|
| **Fruit spine** | **441.7** (highest) |
| Ovary | 114.5 |
| Flower bud | 101.7 |
| Petiole | 100.9 |
| Leaf | 85.9 |
| Male flower | 65.5 |
| Fruit | 58.1 |
| Cotyledon | 57.4 |

**Tau tissue-specificity index: 0.894** — highly tissue-specific, with dominant expression in **fruit spines**. This is consistent with CHS being a key enzyme for flavonoid/anthocyanin pigment accumulation in black cucumber spines.

**Top co-expressed genes** (Pearson r):

| Co-expressed Gene | r | Annotation |
|---|---|---|
| CsaV3_5G010920 | 0.736 | Chalcone-flavonone isomerase (CHI) |
| CsaV3_3G039400 | 0.733 | FSH1 domain-containing protein |
| CsaV3_6G009030 | 0.673 | Flavanone-3-hydroxylase (F3H) |
| CsaV3_7G003260 | 0.655 | Glycosyltransferase |
| CsaV3_7G023390 | 0.650 | Polyol transporter 5-like |

The co-expression with **CHI** and **F3H** — the immediate downstream enzymes in the flavonoid pathway — strongly corroborates its identity as CHS.

**4. Protein Structure**

- **ESMFold pLDDT: 90.7** (Excellent confidence)
- **Predicted binding sites:** Protein-binding interface (56 residues, confidence 0.889) and peptide-binding site (12 residues, confidence 0.718). The protein-binding surface is consistent with CHS functioning as a homodimeric type III polyketide synthase.

💡 You can explore the 3D structure interactively in the **Proteins** module by entering the gene ID CsaV3_3G027830.

**5. Natural Variants**

42 natural variants identified; 7 protein-altering, including:

- **p.E252*** (nonsense, premature stop) — potentially loss-of-function
- 6 missense variants (p.I395L, p.H386D, p.G363R, p.M350T, p.H258Q, p.M50T)

**6. Related Papers (Key Findings)**

1. **Chen et al. (2021),** ***Int. J. Mol. Sci.*** — Identified CHS (Csa3G600020.1) as a key gene in the phenylpropanoid→flavonoid pathway controlling **yellow vs. white cucumber peel**. CHS was downregulated 2.95× in the yellow-peel mutant OL19, and was predicted to drive synthesis of at least three terminal flavonoid metabolites (glycitein, kaempferol, homoeriodictyol).
2. **Li et al. (2020),** ***Horticulture Research*** — Showed that **CsMYB60** directly and indirectly activates structural genes including **CsCHS** to promote flavonol and proanthocyanidin biosynthesis in cucumber black spines. CsMYB60 interacts with CsTATA-box binding protein to regulate CsCHS expression.
3. **Qian et al. (2019),** ***Photochem. Photobiol. Sci.*** — Characterized the expression of all three cucumber **CsCHS** genes under UV-A/UV-B. CsCHS2 displayed a UV-B-enriched expression pattern (pattern 2b) with >30-fold induction, associated with MRE, ACE, and G-box cis-elements in its promoter.
4. **Zhang et al. (2019),** ***Planta*** — The **HEUKCHEEM** gene (B locus) regulates black spine color by inducing anthocyanin biosynthetic genes including CHS. Mutations in HEUKCHEEM abolish CHS transcript accumulation, producing white spines.
5. **Wu et al. (2022),** ***Int. J. Mol. Sci.*** — CHS and CHI were downregulated under low-temperature stress in cucumber, suggesting reduced flavonoid biosynthesis contributes to cold sensitivity; exogenous nitric oxide partially mitigated this.

📚 **References**


1. Chen C, Zhou G, Chen J, Liu X, Lu X, Chen H, Tian Y. Integrated Metabolome and Transcriptome Analysis Unveils Novel Pathway Involved in the Formation of Yellow Peel in Cucumber. *International Journal of Molecular Sciences*, 2021. DOI: 10.3390/ijms22031494
2. Li J, Luan Q, Han J, Zhang C, Liu M, Ren Z. CsMYB60 directly and indirectly activates structural genes to promote the biosynthesis of flavonols and proanthocyanidins in cucumber. *Horticulture Research*, 2020. DOI: 10.1038/s41438-020-0327-z
3. Qian M, Kalbina I, Rosenqvist E, Jansen MAK, Teng Y, Strid A. UV regulates the expression of phenylpropanoid biosynthesis genes in cucumber (Cucumis sativus L.) in an organ and spectrum dependent manner. *Photochemical & Photobiological Sciences*, 2019. DOI: 10.1039/c8pp00480c
4. Zhang C, Win KT, Kim YC, Lee S. Two types of mutations in the HEUKCHEEM gene functioning in cucumber spine color development can be used as signatures for cucumber domestication. *Planta*, 2019. DOI: 10.1007/s00425-019-03244-w
5. Wu P, Kong Q, Bian J, Ahammed GJ, Cui H, Xu W, Yang Z, Cui J, Liu H. Unveiling Molecular Mechanisms of Nitric Oxide-Induced Low-Temperature Tolerance in Cucumber by Transcriptome Profiling. *International Journal of Molecular Sciences*, 2022. DOI: 10.3390/ijms23105615


**Supplemental Figure 7. Full agent session for the question "What is the function, tissue expression profile, and related papers for CsaV3_3G027830?".**

The screenshot shows CucurbitAgent's complete answer, with an expandable work log, numbered evidence sections, in-text citations, and a reference list with DOIs.

**Supplemental Table 1. Module-level data inventory of the CucurbitAgent knowledge foundation**

| Module | Scope / inventory | Key capabilities |
|---|---|---|
| Genes | 9 cultivated species;<br>~256,000 reference genes reconciled across multi-assembly annotations;<br>~9.8M all-vs-all homology hits | Gene annotations; Cross-version ID maps<br>GO/Pfam; ~16.0M natural variants across five species;<br>Orthologs; CDS/protein download |
| Expression | 1,457 RNA-seq samples<br>(species × project × tissue) | Tissue profiles; specificity $\tau$;<br>co-expression neighbors (Top $K = 20$, $\|r\| \geq 0.5$) |
| Proteins | ~242,000 ESMFold models (9 species);<br>~14.2 million GPSite binding-site residues | Gene-resolution fold/confidence;<br>Binding sites (protein/peptide/DNA/RNA/ATP/metal) |
| Literature | 18,305 curated papers;<br>Daily PubMed refresh | Hybrid retrieval (FTS5 + vector + rerank);<br>Passage-level DOI/PMID grounding |

**Supplemental Table 2. Supported languages of the multilingual text interface**

| Language | Native name | Code | Primary regions of use |
| --- | --- | --- | --- |
| English | English | en | Global lingua franca |
| Chinese (Mandarin) | 中文 | zh | China, Singapore, diaspora |
| Hindi | हिन्दी | hi | India, Fiji |
| Spanish | Español | es | Spain, Latin America, US |
| Arabic | العربية | ar | Middle East and North Africa |
| French | Français | fr | France, Canada, W/C Africa |
| Bengali | বাংলা | bn | Bangladesh, eastern India |
| Portuguese | Português | pt | Brazil, Portugal, Lusophone Africa |
| Russian | Русский | ru | Russia, Central Asia |
| Urdu | اردو | ur | Pakistan (RTL) |
| Indonesian | Indonesia | id | Indonesia |
| Swahili | Kiswahili | sw | East Africa |

| Language | Native name | Code | Primary regions of use |
|---|---|---|---|
| German | Deutsch | de | Germany, Austria, Switzerland |
| Japanese | 日本語 | ja | Japan |
| Turkish | Türkçe | tr | Türkiye |
| Vietnamese | Tiếng Việt | vi | Vietnam |
| Korean | 한국어 | ko | Koreas, diaspora |
| Tamil | தமிழ் | ta | India, Sri Lanka, Singapore |
| Italian | Italiano | it | Italy, Switzerland |
| Persian | فارسی | fa | Iran, Afghanistan |
| Hausa | Hausa | ha | West Africa (Nigeria, Niger) |
| Filipino | Filipino | fil | Philippines |
| Malay | Melayu | ms | Malaysia, Brunei |
| Thai | ไทย | th | Thailand |
| Polish | Polski | pl | Poland |
| Ukrainian | Українська | uk | Ukraine |

| Language | Native name | Code | Primary regions of use |
|---|---|---|---|
| Uzbek | O‘zbekcha | uz | Uzbekistan |
| Dutch | Nederlands | nl | Netherlands, Belgium |
| Romanian | Română | ro | Romania, Moldova |
| Greek | Ελληνικά | el | Greece, Cyprus |
| Kazakh | Қазақша | kk | Kazakhstan |
| Swedish | Svenska | sv | Sweden, Finland |
| Uyghur | ئۇيغۇرچە | ug | Xinjiang, China |
| Czech | Čeština | cs | Czechia |
| Bulgarian | Български | bg | Bulgaria |
| Tibetan | བོད་ཡིག | tib | Tibet, China |
| Mongolian | Монгол | mn | Mongolia, China |

**Supplemental Table 3. Data inventory per species across nine cucurbit assemblies**

| Species | Latin name | Reference assembly | Annotated genes | RNA-seq samples (projects) | Natural variants | ESMFold structures | Cross-version ID map |
|---|---|---|---|---|---|---|---|
| Cucumber | Cucumis sativus | Chinese Long v3 | 24,317 | 523 (88) | 5,347,791 | 23,736 | V1↔V2↔V3 |
| Melon | Cucumis melo | DHL92 v4 | 28,299 | 362 (40) | 32,268 | 27,757 | V3.5.1↔V3.6.1↔V4.0 |
| Watermelon | Citrullus lanatus | 97103 v2.5 | 21,917 | 297 (47) | 10,444,284 | 14,901 | V1↔V2↔V2.5 |
| Pumpkin | Cucurbita moschata | Rifu v2 | 30,857 | 62 (15) | 116,322 | 29,898 | V1↔V2 |
| Squash | Cucurbita pepo | MU-CU-16 v4.1 | 27,868 | 94 (11) | 102,721 | 26,813 | single assembly |
| Bitter gourd | Momordica charantia | OHB3-1 (Goya) v2 | 41,016 | 30 (4) | — | 39,839 | single assembly |
| Bottle gourd | Lagenaria siceraria | USVL1VR-Ls v1 | 22,472 | 40 (5) | — | 21,629 | single assembly |

| **Species** | **Latin name** | **Reference assembly** | **Annotated genes** | **RNA-seq samples (projects)** | **Natural variants** | **ESMFold structures** | **Cross-version ID map** |
|---|---|---|---|---|---|---|---|
| Sponge gourd | Luffa cylindrica | P93075 v1 | 31,661 | 21 (8) | — | 30,846 | single assembly |
| Wax gourd | Benincasa hispida | B227 v1 | 27,467 | 28 (4) | — | 26,897 | single assembly |
| **Total** | **—** | **—** | **255,874** | **1,457 (222)** | **16,043,386** | **242,316** | **4 multi-version maps** |

**Supplemental Table 4. CucurbitBench capability coverage matrix (50 items)**

| Capability axis | Benchmark stratum | Items | Question IDs |
|---|---|---|---|
| Cross-version | Genes | 4 | Q02, Q07, Q08, Q09 |
| Gene information | Genes | 9 | Q01, Q03, Q04, Q05, Q06, Q10, Q11, Q12, Q13 |
| Variants | Proteins | 1 | Q26 |
| Expression | Expression | 12 | Q14, Q15, Q16, Q17, Q18, Q19, Q20, Q21, Q22, Q23, Q24, Q25 |
| Protein structure | Proteins | 11 | Q27, Q28, Q29, Q30, Q31, Q32, Q33, Q34, Q35, Q36, Q37 |
| Paper citation | Literature | 8 | Q38, Q39, Q40, Q41, Q42, Q43, Q44, Q45 |
| Cross-module evidence integration | Cross-module integration | 3 | Q46, Q47, Q48 |
| Negative control | Negative control | 2 | Q49, Q50 |
| **Total** | **—** | **50** | **Q01–Q50** |

**Supplemental Table 5. Per-module win rates of CucurbitAgent against general-purpose LLMs on CucurbitBench**

| Capability axis | Items | Pairwise wins / comparisons | Win rate (%) | CucurbitAgent composite | Baseline mean composite |
|---|---|---|---|---|---|
| Cross-version identifier resolution | 4 | 36 / 36 | 100.0 | 93.3 | 26.7 |
| Gene structure & functional annotation | 9 | 81 / 81 | 100.0 | 93.6 | 24.4 |
| Natural-variant interpretation | 1 | 9 / 9 | 100.0 | 94.2 | 23.6 |
| Expression & co-expression | 12 | 108 / 108 | 100.0 | 92.4 | 26.0 |
| Protein structure & binding sites | 11 | 99 / 99 | 100.0 | 87.7 | 27.6 |
| Literature citation tracing | 8 | 72 / 72 | 100.0 | 81.8 | 33.4 |
| Cross-module evidence integration | 3 | 27 / 27 | 100.0 | 94.8 | 24.5 |
| Negative control (out-of-domain) | 2 | 0 / 18 | 0.0 | 69.9 | 83.4 |
| Overall (weighted) | 50 | 432 / 450 | 96.0 | 89.3 | 28.0–30.3 |

**Supplemental Table 6. Evaluation protocol of CucurbitBench and blinded expert review**

| Protocol dimension | Setting |
|---|---|
| Benchmark | CucurbitBench: 50 English items across eight capability axes, balanced across nine cucurbit species |
| Evaluated systems | CucurbitAgent and nine general-purpose LLMs:<br>GPT-4.1-mini, Gemini-2.5-flash, Claude-Haiku-4.5, DeepSeek-V3,<br>Kimi-K2, Qwen-plus, GLM-4.7-flash, Hunyuan-HY3, Seed-2.0-mini |
| Baseline access | A unified API gateway (OpenRouter); no access to the cucurbit databases;<br>same English prompts, single turn |
| Decoding settings | Temperature 0.2; max_tokens 12,000; single turn |
| Gold keys | Exported from the platform's local database and verified by domain experts;<br>no model-generated content used as a scoring basis |
| Raters | Three cucurbit specialists; system identities anonymized (blinded) |
| Scoring dimensions | Factuality, Comprehensiveness, Relevance, Traceability<br>each 0–100 per response |
| Composite score | Factuality 40% + Comprehensiveness 20% + Relevance 20% + Traceability 20%; |

| Protocol dimension | Setting |
| --- | --- |
| | per-item scores averaged over raters |
| Statistics | Pairwise wins per item against each baseline (450 comparisons, 0 ties);<br>overall win rate with 95% CI by bootstrap (10,000 resamples);<br>Wilcoxon signed-rank test per baseline (all $P < 0.001$);<br>inter-rater alpha reported per scoring round |
| Collection | Final benchmark collection 2026-08-25/26 (all 10 systems × 50 items;<br>CucurbitAgent answers re-collected with full reference lists) |
